# High resolution shotgun metagenomics: the more data, the better?

**DOI:** 10.1101/2022.04.19.488797

**Authors:** Julien Tremblay, Lars Schreiber, Charles W Greer

## Abstract

In shotgun metagenomics (SM), the state of the art bioinformatic workflows are referred to as high resolution shotgun metagenomics (HRSM) and require intensive computing and disk storage resources. While the increase in data output of the latest iteration of high throughput DNA sequencing systems can allow for unprecedented sequencing depth at a minimal cost, adjustments in HRSM workflows will be needed to properly process these ever-increasing sequence datasets. One potential adaptation is to generate so-called shallow SM datasets that contain fewer sequencing data per sample as compared to the more classic high coverage sequencing. While shallow sequencing is a promising avenue for SM data analysis, detailed benchmarks using real data are lacking. In this case study, we took four public SM datasets, one massive and the others moderate in size and subsampled each dataset at various levels to mimic shallow sequencing datasets of various sequencing depths. Our results suggest that shallow SM sequencing is a viable avenue to obtain sound results regarding microbial community structures and that high depth sequencing does not bring additional elements for ecological interpretation. More specifically, results obtained by subsampling as little as 0.5M sequencing clusters per sample were similar to the results obtained with the largest subsampled dataset for the human gut and agricultural soil datasets. For the Antarctic dataset, which contained only a few samples, 4M sequencing clusters per sample was found to generate comparable results to the full dataset. One area where ultra-deep sequencing and maximizing the usage of all data was undeniably beneficial was in the generation of metagenome-assembled genomes (MAGs).

**Key points:**

– Three public multi-sample shotgun metagenomic NovaSeq datasets totalling 12,389,583 and 202 Gb, respectively were analyzed at various sequencing depths to evaluate the accuracy of shallow shotgun metagenomic sequencing using a high resolution shotgun metagenomic bioinformatic workflow. A synthetic mock community of 20 bacterial genomes was also analyzed for validation purposes.
– Datasets subsampled to low sequencing depths gave nearly identical ecological patterns (taxonomic and functional composition and beta-alpha-diversity) compared to high depth subsampled datasets.
– Rare taxa and functions could be uncovered with high sequencing depth vs. low sequencing depth datasets, but did not affect global ecological patterns.
– High sequencing depth was positively correlated with both quantity and quality of recovered metagenome-assembled genomes.

## Introduction

DNA sequencing costs have decreased dramatically in recent years. With the introduction of Illumina’s short reads-based NovaSeq system, the cost of sequencing has reached a new low mark. In the field of metagenomics, it is challenging to estimate the required output of the generated sequences needed to get a satisfactory level of coverage and is particularly true for complex environments, like soil and animal guts. To date, sequencing technology could hardly reach the saturation of such complex environments and common wisdom in estimating sequencing output for such environments was simply to generate the largest possible amount of sequence within budget constraints. However, the amount of data generated by the latest iteration of sequencing systems (i.e. Illumina’s NovaSeq; Oxford Nanopores’s Promethion) has reached a point where it can now far exceed computational capacity in shotgun metagenomics (SM) analysis workflows [1] like what is loosely referred to as high-resolution shotgun metagenomics (HRSM). In this type of workflow, raw sequence libraries are usually controlled for quality, trimmed and *de novo* co-assembled. Quality controlled reads are then mapped back to the co-assembly in order to estimate contigs and gene abundance to ultimately generate abundance matrices and metagenome-assembled genomes (MAGs) [2]. The most critical aspect of this type of workflow is probably the *de novo* co-assembly of all the relevant sequence libraries generated for a given project. This step ideally requires what is referred to as a large compute node: a computer node usually equipped with tens of cores and large amounts of Random Access Memory (RAM) to adequately process all short reads into a data structure to perform a *de novo* co-assembly of many samples at once. While some *de novo* assembly software are written to handle multiple compute nodes (i.e. MetaHipMer [3], Ray Meta [4]) through distributed-memory systems paradigms such as Massively Parallel Interface (MPI), they are not the most practical to implement and use because of their inherent configuration complexity and do not necessarily generate the most accurate assemblies [5]. Moreover, the most widely used and arguably some of the best *de novo* assembly software are written as a single node solution (for instance metaSPAdes [6,7], MEGAHIT [6]). The question of what assembling software package is the most performant is currently the subject of debate [8] and is beyond the scope of this study. Although the analysis pipeline used for this study supports using metaSPAdes, we ended up using MEGAHIT for our analyses because it was the only viable option to process this objectively large dataset in an acceptable amount of time. It is to be noted that in order to circumvent the issue of having enough RAM resource to perform a large multi-sample co-assembly, some workflows instead favor performing *de novo* assemblies for each sample separately (for instance, see [9]). This approach, however, prevents the optimal analysis of end results as it generates significant redundancy in assembled contigs and MAGs, making it intractable to directly compare abundance between samples. In contrast, co-assembling all samples together has the advantage of creating one single reliable reference baseline on which all samples can be easily compared/analyzed and increases the power of segregating contigs during MAGs generation.

Here we investigated the end results of a typical HRSM workflow using the largest public Illumina Novaseq6000 dataset available at the moment of writing (PRJNA588513 [10]). This dataset holds 912 paired-end (2 x 150 bp) human gut microbiome shotgun metagenomic sequence libraries representing 12,389 giga-bases (Gb) of raw sequence data for a total of 5.6 terabytes (TB) of compressed fastqs. Co-assembling an amount of sequence data of this magnitude is not achievable on the vast majority of existing large compute node hardware as it would require unrealistic - or at least hardly accessible - amounts of both RAM and compute time. To circumvent this limitation, we emulated various level of shallow sequencing by iteratively randomly subsampling this sequence data to up to a total of 2,927 Gb (3.2 Gb / library) which saturated our largest compute node (i.e. one ‘4 x Intel Xeon E7-8860v3 @ 2.20GHz; 3 TB RAM; 64 cores’ node) and dissected the end results (taxonomic profiles, alpha-beta-diversity and MAGs). We performed the same exercise with additional more modest datasets consisting of 18 NovaSeq6000 libraries (2 x 150 bp; 583 Gb) from samples obtained from Antarctic soil environments (PRJNA513362 [9]), 72 HiSeq2500 libraries (2 x 125 bp; 202 Gb) obtained from an agricultural soil [11] and 6 HiSeq2500 libraries (2 x 125 bp; 8 Gb) obtained from commercial mock communities (PRJNA873699; [12]).

## Results

### Experimental design

A large-scale shotgun metagenomic sequencing project consisting of 912 gut microbiome samples collected from individuals across six provinces of China was recently published [10]. These samples were sequenced on numerous lanes of a NovaSeq6000 system and yielded a total of 12,232 Gb representing 5.7 TB of compressed fastqs. To our knowledge, this dataset is one of, if not the largest SM dataset to be made publicly available as part of a single project and provides an opportunity to determine if generating that much data was necessary to obtain meaningful results or validate hypotheses. As sequencing systems keep getting more performant in generating data, this becomes an increasingly important question as processing 5.7 TB of raw SM data is not a trivial endeavour and generates countless intermediate files, inflating the storage and compute resources requirements to properly analyze the end results. This dataset is therefore unsuitable for a HRSM workflow as-is (*i.e*. without subsampling or reducing the raw reads input load). With the current trends in sequencing technology development, it is not unreasonable to expect a similar amount of sequencing data being routinely generated in the near future for any given SM project.

We therefore downloaded the raw fastq files related to this project and processed them with an iterative subsampling strategy in order to determine if smaller subsets of this dataset would be sufficient to reach sound biological conclusions. Like the vast majority of research units, we did not have a compute node with enough RAM to perform a *de novo* co-assembly for the entirety of such a large dataset. The best resources we had access to at the moment of writing were 3 TB RAM 64 core nodes. Therefore, for each dataset, we adopted a strategy (Fig. 1) in which we performed quality control of all the raw sequence data files to trim adapters and remove sequencing artefacts and contaminants which yielded what we will refer to as quality controlled reads. We iteratively subsampled the quality-controlled fastqs to obtain 0.1, 0.5, 1, 4, 8 and 12 million of sequencing clusters (*i.e.* one sequencing cluster represents two reads of 150 bp each) for each sample which amounts to approximately 0.03, 0.14, 0.27, 1.07, 2.14 and 3.21 Gb per library, respectively (Table I). For each of these subsampled pools of libraries, what we refer to a standard HRSM workflow (*i.e.* standard workflow in Fig. 1) was executed and essentially achieved a *de novo* co-assembly, mapping of quality controlled reads on the co-assembly to generate contigs and gene abundance matrices, computation of alpha and beta diversity, assignation of both taxonomic lineages and KEGG orthologs (KO) and MAGs generation. In parallel, the all reads mapped (arm) workflow was executed. This workflow consists of mapping the complete full size fastq libraries (*i.e.* In the context of this article, a sequencing library is considered analogous to a sample) to estimate read abundance, instead of using only a fraction of the reads like it is the case in the standard workflow (Fig. 1). Ultimately, the only difference between the standard and arm workflow for any given subsampled dataset, is the number of reads that were mapped on the co-assembled reference (Fig. 1): in the standard workflow, only the subsampled libraries are mapped to the co-assembly to estimate contig and gene abundance, whereas in the arm workflow, the full size (not subsampled) libraries are mapped against the co-assembly. The arm workflow’s purpose is to allow using the maximum amount of sequencing data should the compute node used to perform the co-assembly does not have enough RAM to process the whole sequence dataset. We favored the MEGAHIT software to perform the *de novo* co-assembly because of its speed, its capacity to handle relatively large sequencing datasets and because it is designed to be executed on a single compute node. We performed similar *in silico* experiments with three additional SM sequence dataset of lower magnitude obtained from natural extreme Antarctic environments (523.16 Gb; 18 samples; PRJNA513362 [9]), agricultural soil samples (202.09 Gb; 72 samples; PRJNA448773 [11]) and a microbial mock community (PRJNA873699; [12]). Although these three latter datasets are not as massive as the one described above, they are nonetheless substantial and typical of recent SM datasets, except for the mock community which was included for validation purposes.

**Figure 1.**
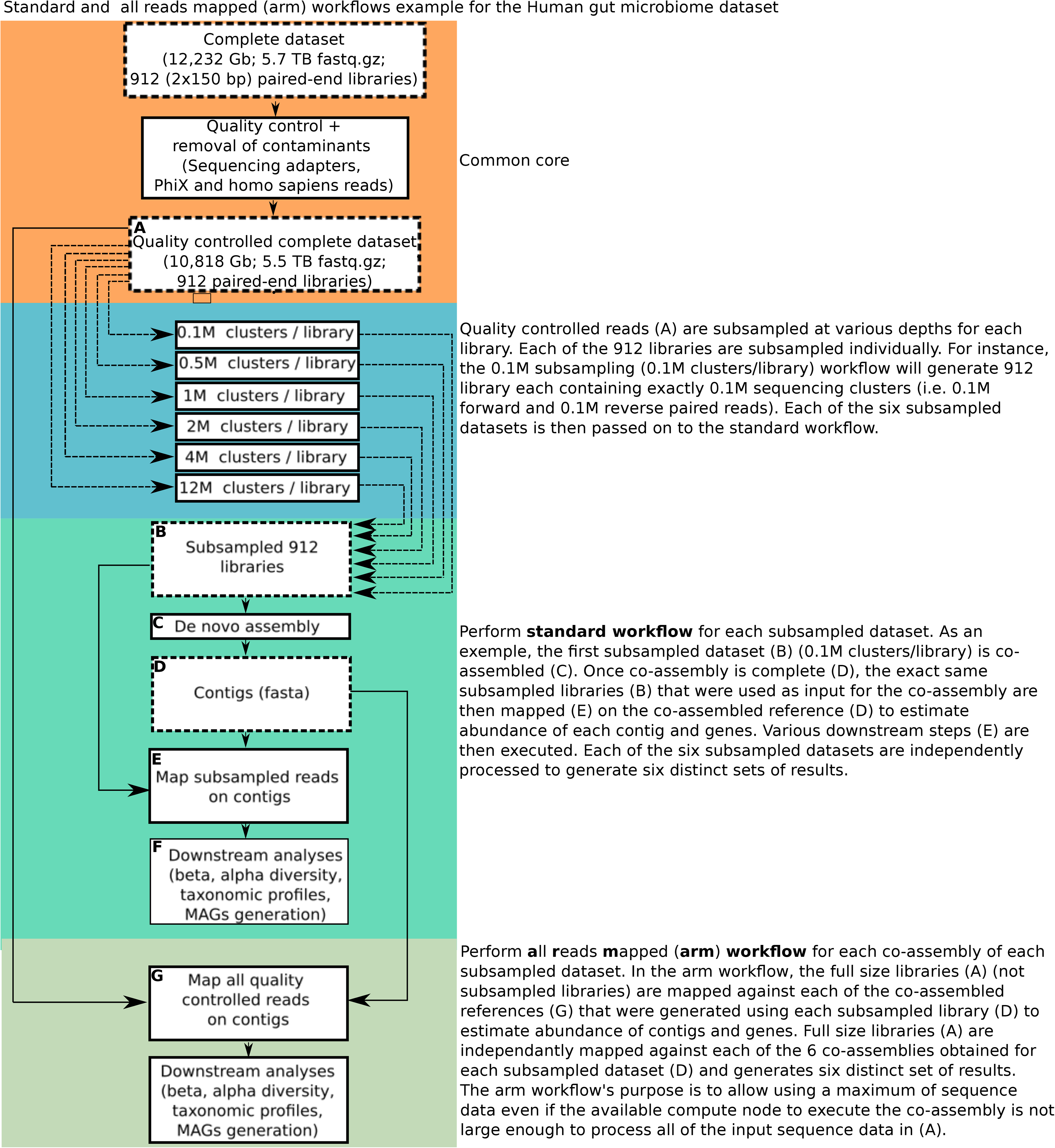
Experimental design used to investigate the effects of the number of reads input on the end results of each shotgun metagenomics dataset investigated here. For each dataset, a common core workflow was performed where each library was trimmed according to their quality profile and filtered for common contaminants. Quality controlled libraries were subsampled at 0.1, 0.5, 1, 4, 8 and 12 million clusters (*i.e.* times two for the number of reads). Subsampled quality controlled libraries were then co-assembled and analyzed for downstream analyses for the standard workflow. In the ‘all reads mapped’ (arm) workflow, all of the quality controlled reads were mapped against their corresponding co-assembly to estimate contig and gene abundance and investigate how it affects end results. The Antarctic, agricultural soil and mock communities datasets were small enough so that they could be co-assembled and processed at once.

### The amount of resources required for de novo assembly increases almost linearly with the number of input reads used for co-assembly

From Table I and Figure S1, we show that the amount of consumed RAM increases linearly with the number of input reads fed into the assembler, which suggests that the internal memory data structure of kmers still does not saturate as we reach the maximum memory capacity of the compute node during the assembly process. Interestingly, this holds true for the simple mock community dataset as well for which kmer saturation is not reached, despite its high per genome sequencing depth. Given the near perfect linear correlation between the amount of bp and consumed RAM, significantly more memory would have been required to take into account all of the sequencing data of the human gut dataset (12 Tb), which we estimated to be approximately 10 TB of RAM (Table I).

### De novo co-assembly contiguity is positively correlated with the number of input sequences used for co-assembly

For all four datasets, there is a clear linear correlation between the quantity of bases used to perform the co-assembly and the contiguity of the generated assembly (Table I; Fig. S1). Simply put, the more the reads fed into the co-assembly, the more and longer contigs are obtained. The number of quality controlled reads mapping on the co-assembly is a metric that can inform on the quality of the assembly. For the human gut dataset, the percentages of properly aligned reads were somewhat lower for the 0.1M cluster / library workflow (Fig. S2) while inputs from 0.5M onward showed an average mapped reads rate of approximately 90%. The number of reads mapped from the arm workflows - that is, mapping all the quality controlled reads from the dataset on the subsampled co-assemblies (see experimental design in Fig. 1) - showed a high mapping rate (> 90%) suggesting that while only a subset of reads were used to generate the co-assemblies, the generated contigs managed to catch the vast majority of the complete full size libraries. A similar trend is observed for the other and smaller datasets (Antarctic soil, agricultural soil and mock communities), but with the saturation inflection point being generally reached at higher levels of subsampling.

### Alpha and beta- diversity

Beta diversity (Bray-Curtis dissimilarity index) comparison between various co-assemblies suggests that the amount of input of reads in the co-assembly does affect the overall population structure (Fig. 2). The Spearman correlations (Mantel tests between Bray-Curtis dissimilarity matrices) between low (0.1M, 0.5M and 1M clusters) and high (4M, 8M and 12M (and complete dataset for the Antarctic, agricultural soil and mock communities datasets)) input configurations were consistently showing lower values compared to high vs high configurations. The contigs and gene richness diversity index were found to be good indicators of each participant’s microbiota composition and is visually highlighted in the PCoAs (Fig. S3A) in which participants cluster according to their diversity quintiles in essentially identical patterns across all workflow configurations. For the Antarctic dataset, the workflow configuration that showed the lowest correlations with the others was the 0.1M - arm design with Spearman *r* statistics values of approximately 0.85. Accordingly, the PCoAs of this dataset gave nearly identical patterns for all workflows except the 0.1M - arm design which is significantly different (Fig. S3B). The agricultural soil dataset had low Spearman correlation values between 0.1M and 0.1M - arm vs other configurations (Fig. 2C), but correlations were however much higher for the slightly more subsampled 0.5 and 0.5 - arm datasets compared with higher input configurations (Fig. 2C and Fig. S3C). The only exception to this is for the mock community where the 0.1M datasets show high correlation to high input configurations (Fig. 2D).

**Figure 2.**
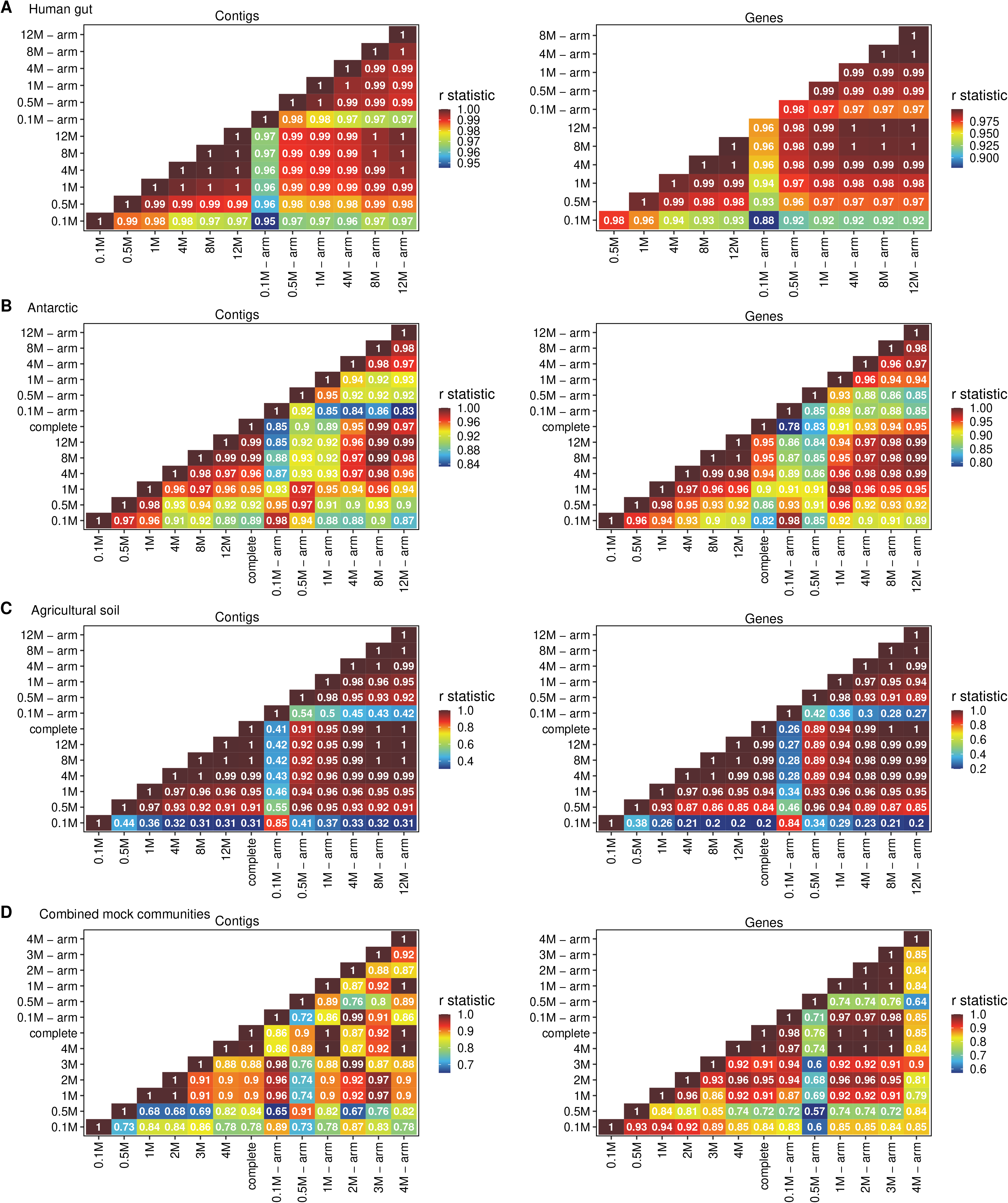
Heatmaps of Spearman correlation coefficients (Mantel test) of normalized (TMM method from edgeR) contig and gene abundance Bray-Curtis (beta diversity) dissimilarity matrices between each workflow for the A) human gut, B) Antarctic, C) agricultural soil and C) mock communities datasets. Higher coefficient means higher correlation between distance matrices.

Observed contigs and observed genes indexes as a function of sequencing efforts (Fig. 3) suggest that saturation is reached at the 4M subsampled clusters for the human gut dataset (Fig. 3A) while both metrics are still in the exponential phase for the Antarctic dataset (Fig. 3B) and agricultural soil (Fig. 3C). The only exception to this is the observed contigs of the mock communities (Fig. 3D) for which the number of contigs decreases with high input workflows.

**Figure 3.**
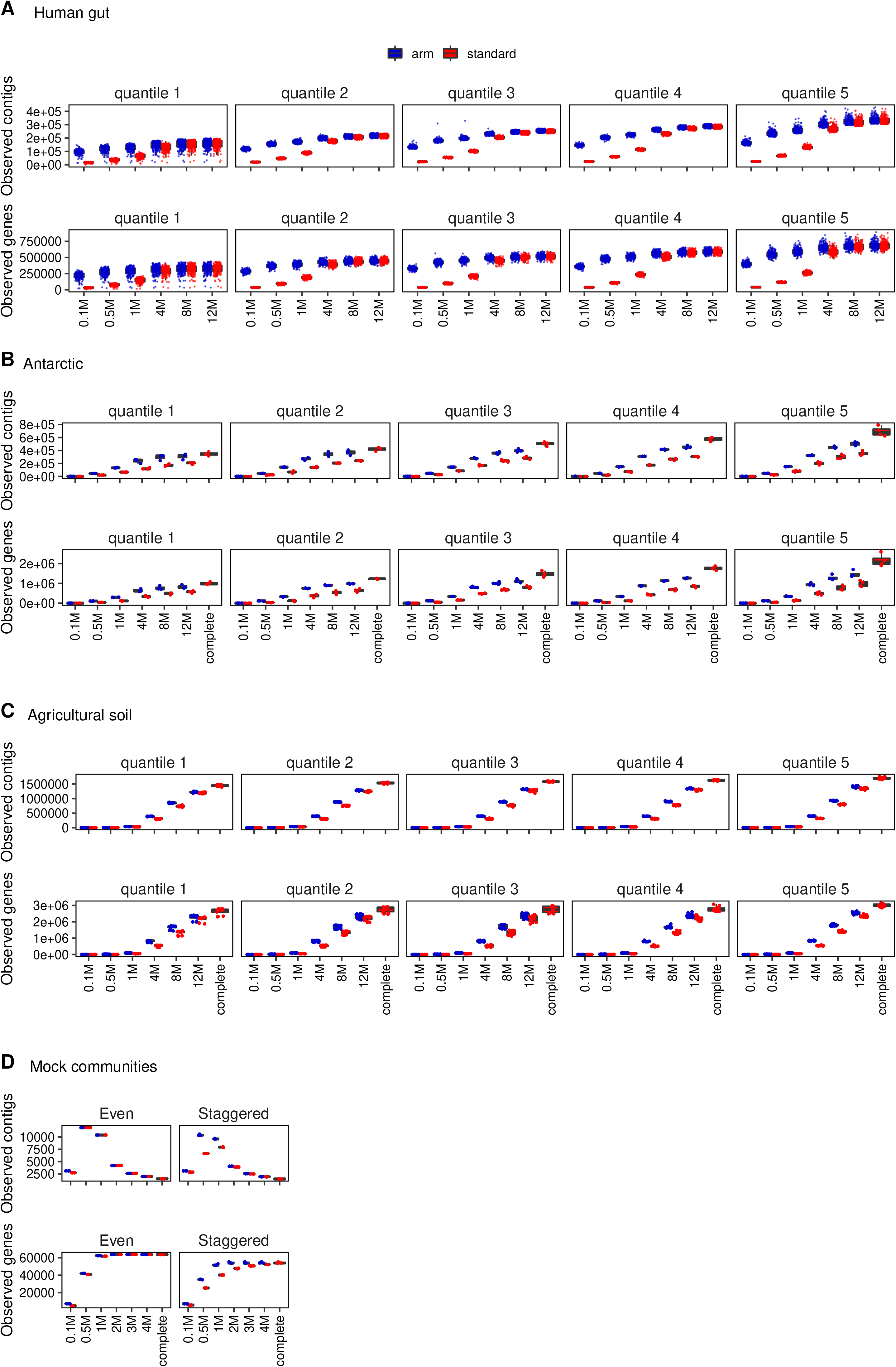
Observed contigs and genes indices binned by quintiles for the A) human gut and B) Antarctic datasets. To improve ease of data visualization, each data point was binned in a quintile. The data points for a given quintile correspond to the same data points from one boxplot to another. Diversity indices were computed from raw contigs or genes abundance tables using RTK v0.93.2.

### Taxonomy and KO

Correlation at the taxonomic and functional level was also assessed and followed similar trends to what was found in the beta diversity comparisons of Figure 2 with low read inputs showing lower correlation against high read input configurations (Fig S4). Even though workflows with more input sequences are associated with a higher number of recovered taxa (Fig. S4 - barplots and Venn diagrams), these “rare” taxa only account for a minor fraction of the total reads (Fig. S4 lower right panels) and the quasi-totality of reads are associated with taxa common to low input (0.1M, 0.5M) workflows. For each dataset, relative abundance profiles of a selection of some of the most overall abundant taxa (Fig. 4) and KOs (Fig. 5) were generated to further validate their consistency across all workflows. For the human gut dataset, taxa abundance generally correlates between data input loads, except for contigs assigned to the genus *Prevotella* that show significant variation in the 0.1M, 0.5M and 1M cluster workflows (Fig. 4A). Interestingly, the same sequencing cluster input loads processed with the arm method correct these variations and bring them to the same stability level as the other workflow configurations. To confirm that *Prevotella* abundance variation in the 0.1M, 0.5M and 1M standard workflows was not a bias caused by the random nature of library subsampling, these workflows were repeated three independent times each with a different seed value during the random subsampling step. Resulting taxonomic profiles show that *Prevotella* still shows appreciable variation in these low-input workflows (Fig. S6) and that no biases were introduced by the subsampling process. Important ecological trends like the overrepresentation of *Coproccus* and prevalence of *Bacteroides* and *Megamonas* in participants harboring a high diversity microbiota are consistent across all workflows. Except for the 0.1M clusters workflow, functional profiles were in general similar between other input workflows (Fig. S5). For the human gut dataset, significantly differentially abundant KOs were determined between low diversity (quintile #1) and high diversity (quintile #5) groups and their abundance profiles were similar between workflows as shown by the selected KOs K01593, K11444, K18143, K22225, K22607 that were found to be more abundant in low diversity participants and K00178, K16149 that were more abundant in high diversity participants. As shown in Fig. 5A, the abundance of these KOs are nearly identical for all participants in all workflows.

**Figure 4.**
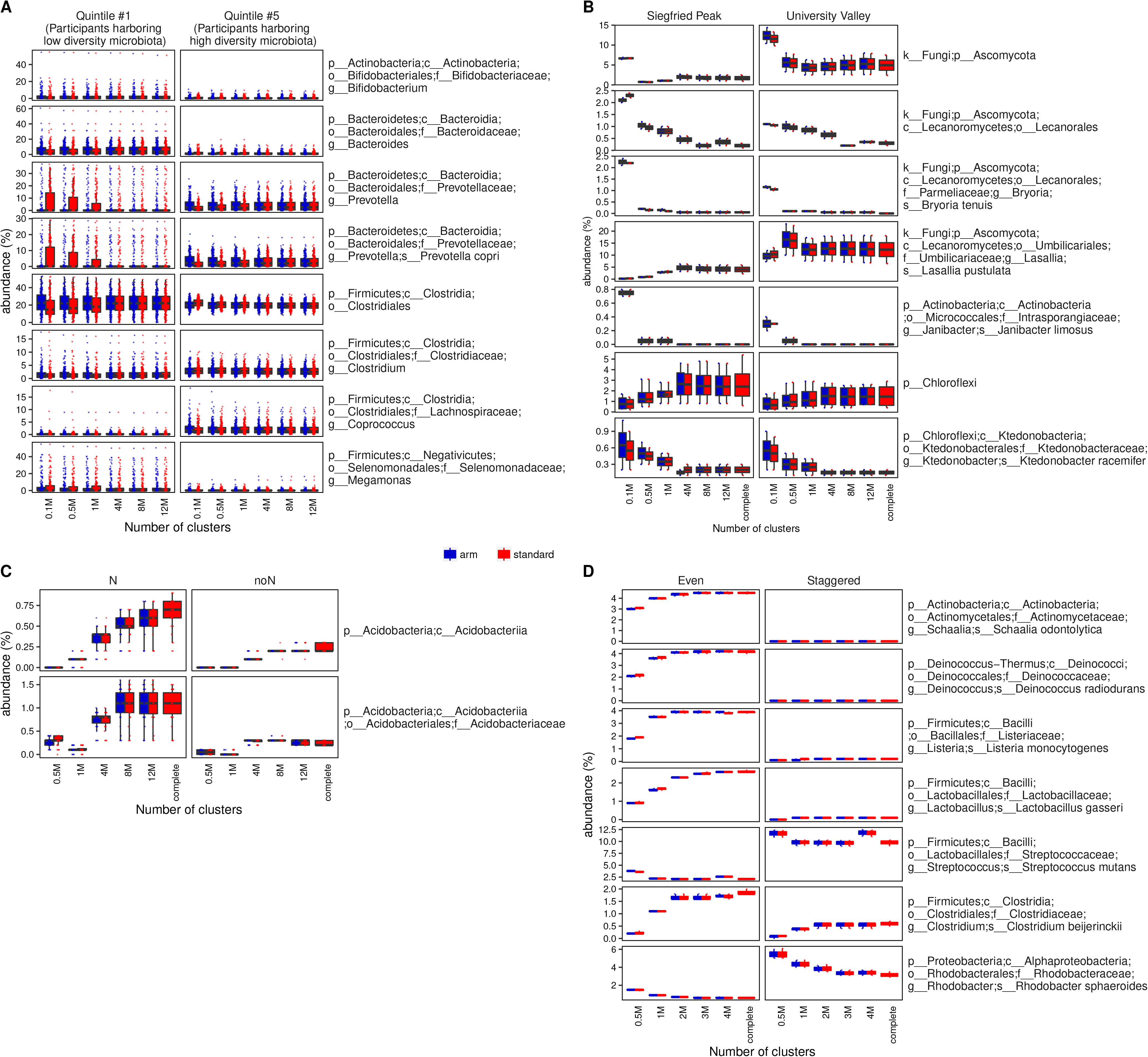
Relative abundance profiles of selected taxa for the A) human gut, B) Antarctic, C) agricultural soil and D) mock communities datasets for each of the described workflows. A) Selected taxa are shown for quintiles #1 (participants with low diversity microbiota) and #5 (participants with high diversity microbiota). B) Selected taxa are shown for the Siegfried Peak and University Valley locations. C) Selected taxa are shown for the nitrogen-rich samples vs samples that were not fertilized with nitrogen. D) Selected taxa for the mock communities.

**Figure 5.**
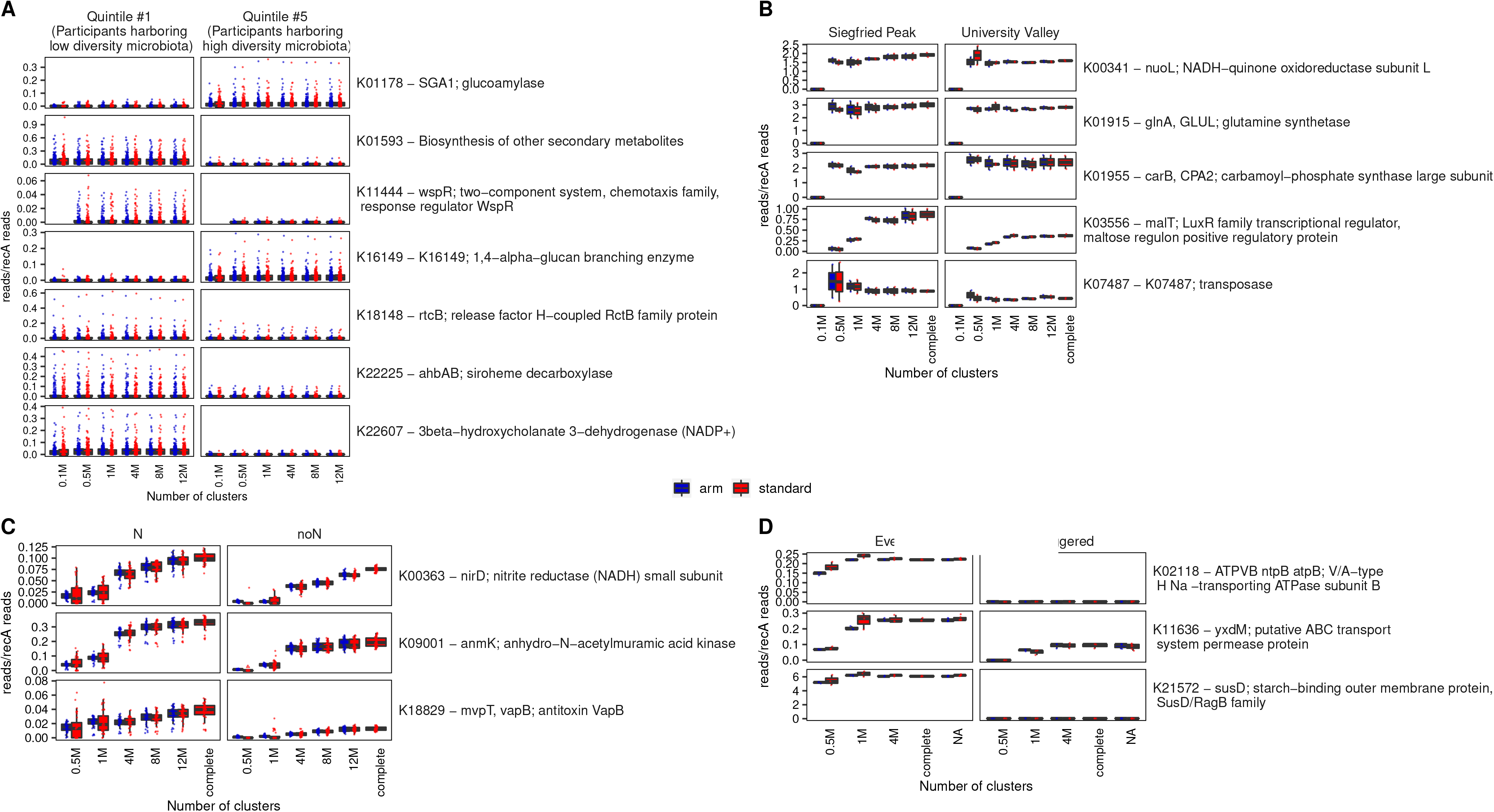
Functional abundance profiles of selected KOs for the A) human gut, B) Antarctic, C) agricultural soil and D) mock communities datasets for each of the described workflows. B) Abundance profiles of a selection of significantly differentially abundant KOs for each workflow for quintiles #1 and #5. KOs were identified by using a one-way anova between samples of quintiles #1 and #5 followed by a Bonferroni correction. KOs having a corrected p-value < 0.01 and a 8 times fold-change between the two quintiles were selected. D) Abundance profile of selected KOs for these same two locations. The Antarctic dataset was unreplicated and was therefore not conducive for feature selection by a statistical text.

For the Antarctic dataset, which contained 18 samples only, a similar trend is observed with the relative abundance of selected taxa higher in low input workflows (0.1M, 0.5M and 1M), but with no correction by the arm method (Fig. 4B). This is also observed for KOs relative abundance as illustrated by the selected KOs relative abundance through all workflows in Fig. 5B. In this dataset, ecological trends are consistent across workflows that contain 4M sequencing clusters or more, but lower input workflows show taxonomic and functional abundance profiles significantly different from the complete dataset. For the agricultural dataset, two taxa (*Acidobacteriia* and *Acidobacteriaceae*) found to be more abundant in fields fertilized with nitrogen were plotted across workflow configurations and their abundances are consistent across sequencing depths. Taxa and KOs abundance between staggered and even mock communities was also examined and showed expected abundance matching with their spiked-in concentrations (Fig 4D; Fig 5D).

### MAGs

The relationship between sequencing input and MAGs generation yield and quality was also investigated and showed that the yield, % contamination and completeness are positively correlated with the amount of input sequences in each workflow of each dataset (Fig. 6). Moreover, the arm workflows consistently generated higher yield MAGs of better quality compared to the standard workflows.

**Figure 6.**
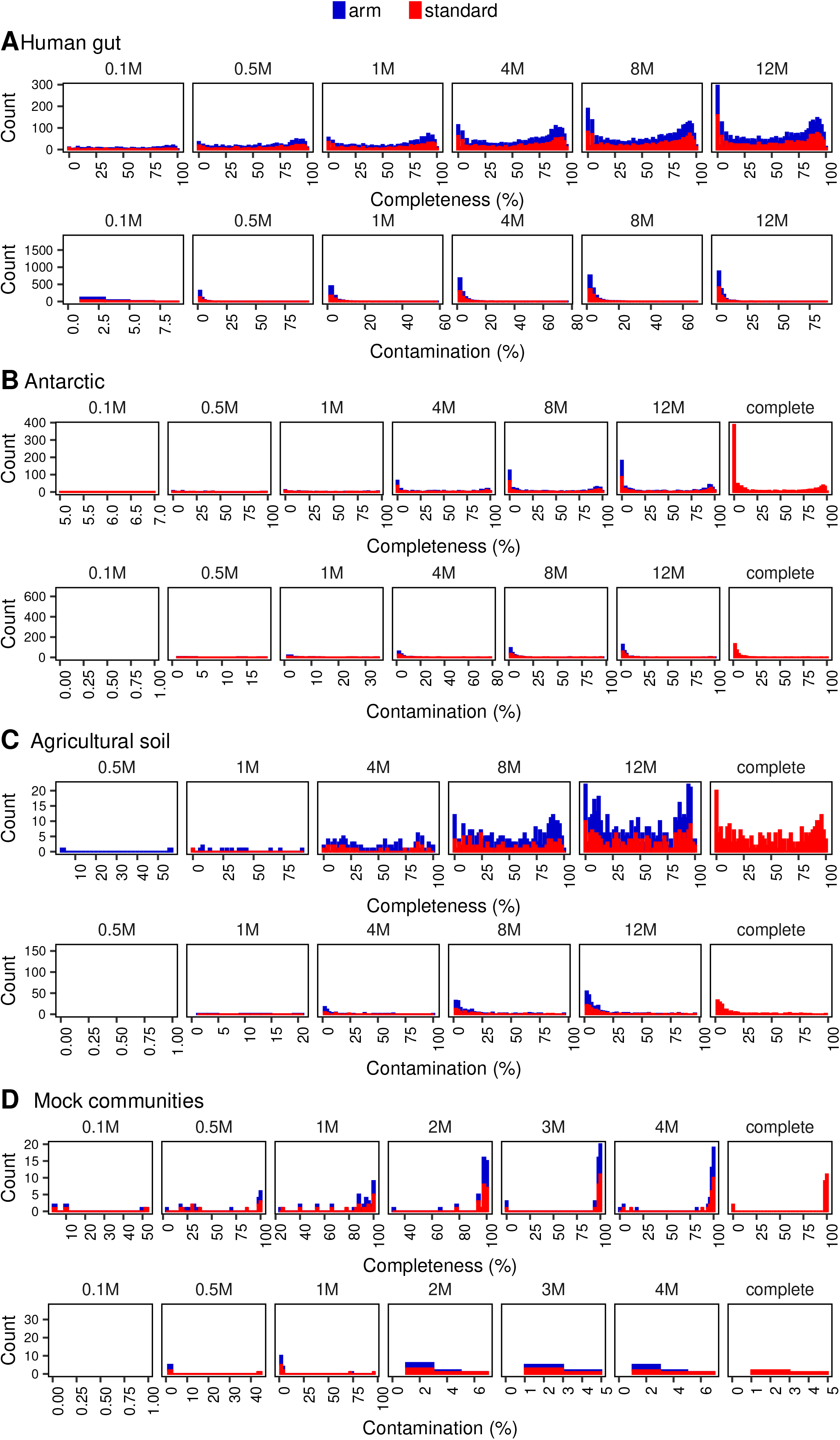
Histograms of MAGs quality assessment metrics for each workflow of the A) human gut and B) Antarctic, C) agricultural soil and D) mock communities datasets. MAGs were generated with MetaBAT2 v2.12.1 and Completion % and contamination % were obtained with CheckM v1.1.3.

### rpoB-based analysis to estimate species and community coverage

The single-copy marker gene *rpoB* [13] was used as a proxy for the detection and abundance of community members, with each unique *rpoB* gene assumed to represent one species. Based on this approach, the communities of the human gut and the Antarctic soil datasets represent communities of low species richness, while the agricultural soil dataset represents communities of high species richness (Fig. 7A). At the same time, communities of the agricultural soil dataset also featured highest community evenness J [14], while communities of the human gut and the Antarctic represent communities of lower evenness (Fig. 7C). Using the complete dataset of each sample as a reference, we estimated that at least 95% of the communities of the human gut and Antarctic datasets would have already been covered [for details on the community coverage concept see Chao & Jost, 2012 [15] by the 4M subsets (Fig. 8B). For the agricultural dataset this threshold would have only been achieved by the 8M subset. In contrast, for all datasets the majority of the species richness (95%) was only covered by complete datasets (Fig. 8A). A similar *rpoB* analysis carried out with the mock community dataset, showed that the majority (>99%) of the even and staggered mock communities were already covered by the 0.5M subsets. The corresponding species coverage analysis, showed that for the staggered community only 12 out of 20 could be detected even when using the complete dataset (*ca*. 0.69 Gbp per sample). On the other hand, the subsets even mock community dataset consistently overestimated the real species richness. Here, the real richness value of 20 species was only obtained again using the full dataset (ca. 0.58 Gbp per sample). We hypothesize that this is a result of the inability of the sequence assembler to generate full-length, contiguous *rpoB* gene assemblies at lower sequencing depths.

**Figure 7.**
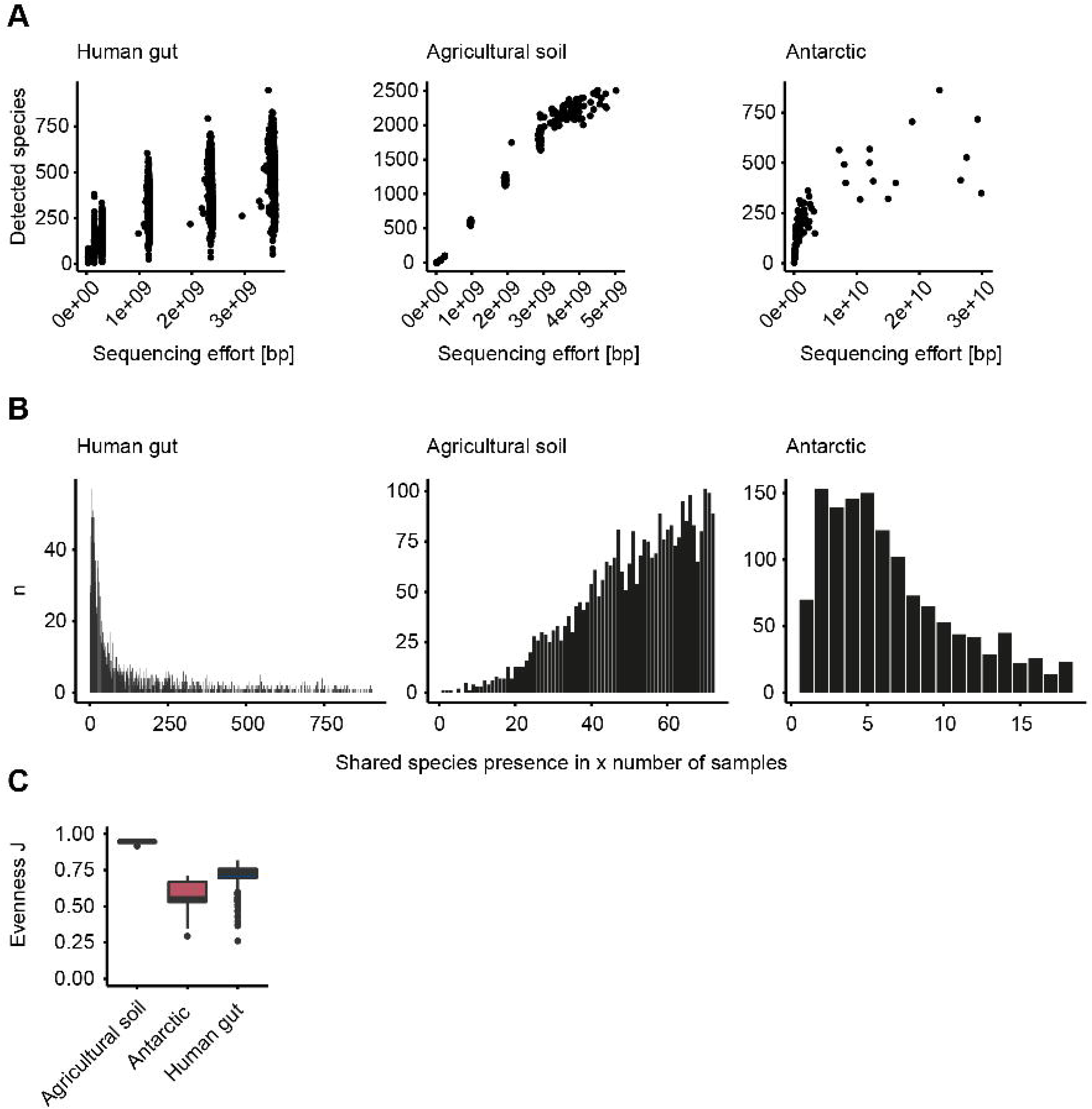
General characteristics of the metagenomic datasets based on the single-copy marker gene *rpoB*. (A) The number of detected species as a function of sequencing effort. The sequencing effort was corrected by apparent contamination of each dataset, so that the shown sequencing effort represents the true sequencing effort of the microbial component of the dataset. (B) Histograms of the frequency [n] of species shared between multiple samples of the same dataset. (C) A summary of the community evenness of the datasets expressed in terms of Pielou’s evenness index J. All metrics were calculated assuming that each unique *rpoB* gene represents one species.

**Figure 8.**
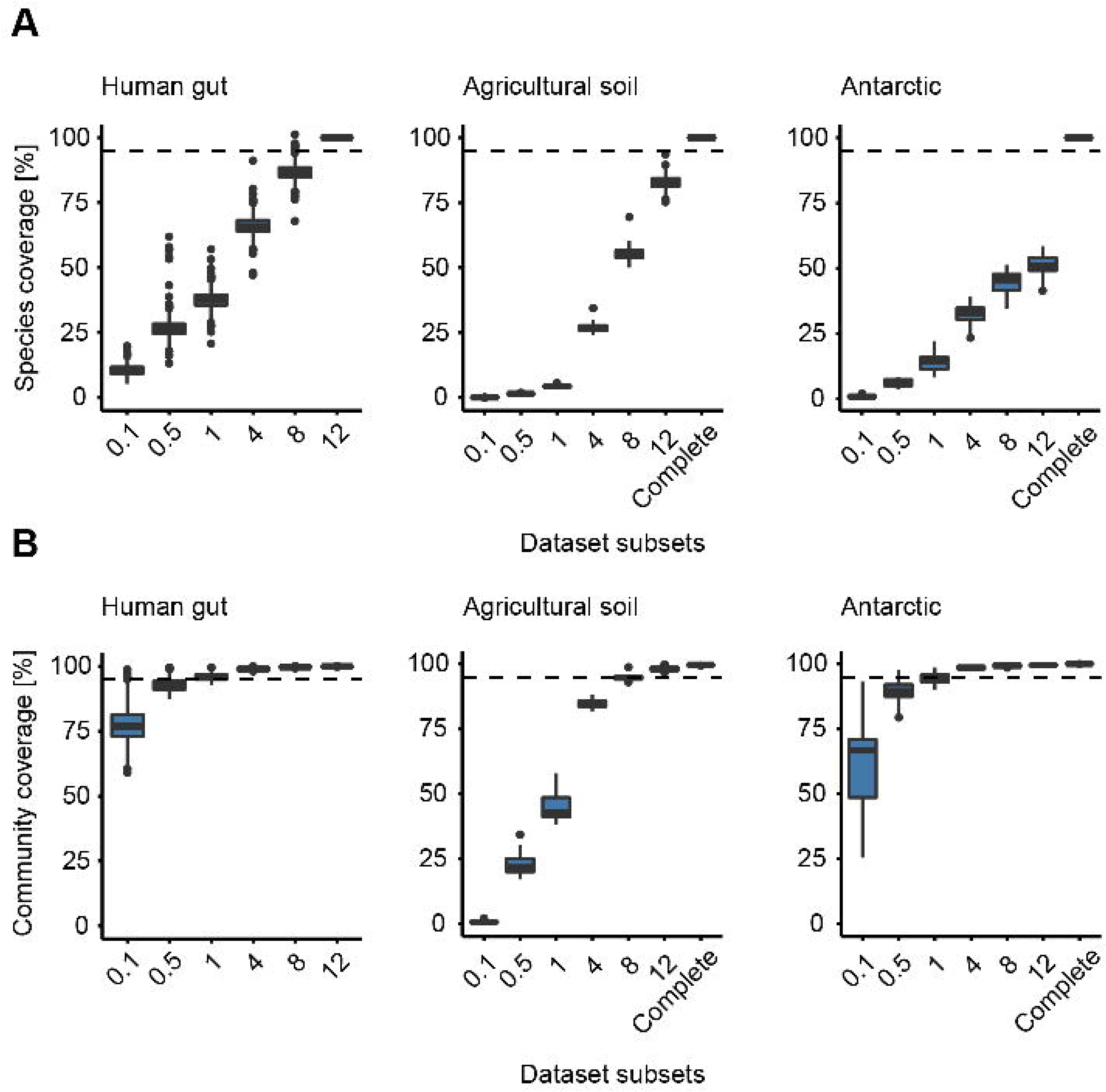
Community representation of dataset subsets based on the *rpoB* gene. (A) Relative species coverage as a function of sequencing depth. This metric was calculated on a by sample basis and by relating the number of detected species of a subset to the overall species number in the complete dataset. (B) Community coverage [Chao & Jost, 2012] as a function of sequencing depth. Assuming the complete dataset of each sample represents the true community, these boxplots show how well the subsets would have covered this true community. The dashed lines indicate 95% species or community coverage, respectively.

### The effect of additional samples

Using the *rpoB* data, we also simulated how community and species coverage would change as a function of the number of sequenced samples under the constraint of a constant sequencing effort. For this analysis, we defined the overall meta-community of each dataset as a reference. The simulations showed that the achieved community and species coverage increased with the number of sequenced samples. Even in the extreme case where all samples of a dataset would have been sequenced together (very shallowly) to the depth of a single original sample, a very high coverage (>95%) of the meta-community would have been obtained (Fig. 9C). At the same time, more shallow sequencing appears to clearly limit the species coverage that can be maximally achieved. We hypothesize that this limit is primarily affected by the proportion of species that are only shared by very few samples. Consistent with this, we observed the highest maximal species coverage (87%) of the simulations for the agricultural soil dataset, where the majority of species appears to be ubiquitously present in the majority of samples (Fig. 7B). In contrast, this maximal species coverage was only 43% and 60% in the human gut and the Antarctic datasets, respectively, which feature a high proportion of species that were only detected in a small number of samples (Fig. 7B).

**Figure 9.**
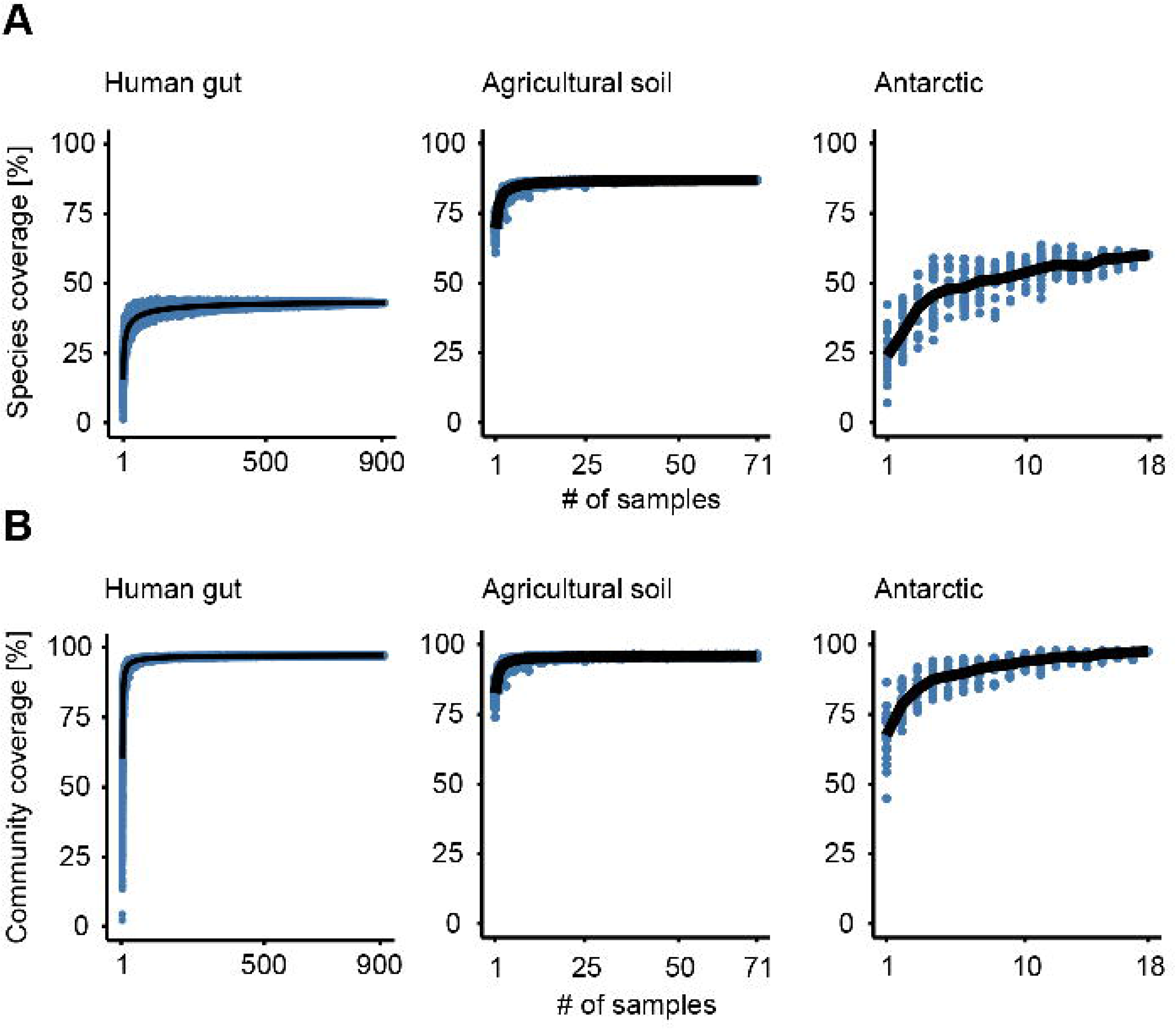
Species and community coverage as a function of the number of samples. This figure shows the simulated effects of adding additional samples while maintaining a constant overall sequencing depth on the species and community coverage. The coverage values were calculated using the combined meta-community of each dataset as a reference.

**Figure 10.**
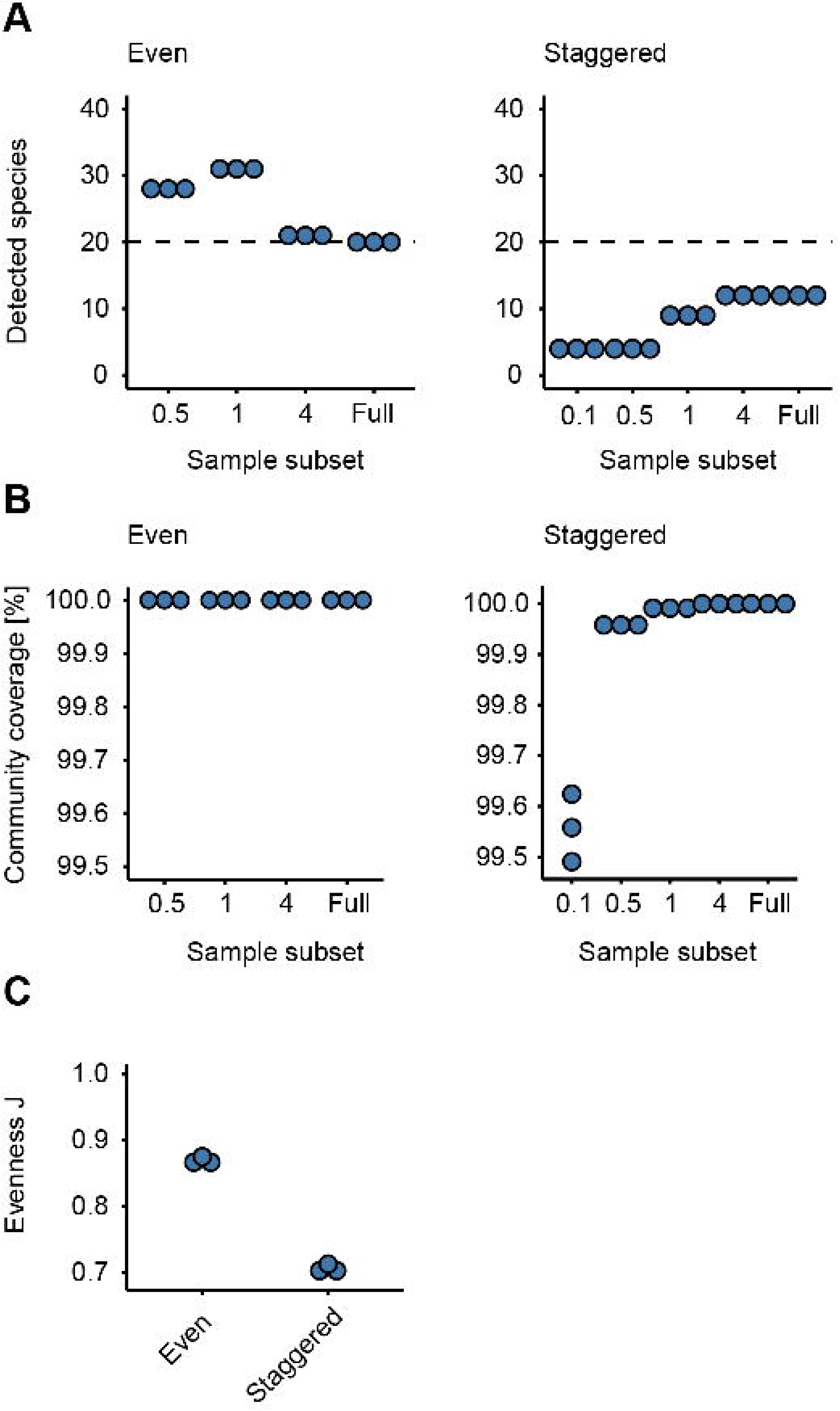
Species discovery (A) and community coverage (B) of the even and staggered mock communities as a function of sequencing depth. (C) Community evenness of the even and staggered mock communities based on the total (full) sequencing depth achieved for each sample.

## Discussion

### Shallow sequencing vs high depth sequencing

Shallow shotgun metagenomic (SSM) sequencing has been the subject of only a few studies so far [16–18] and focused mainly on the correlation between SSM and 16S rRNA amplicon sequencing. They were done with artificial datasets or executed with reads-based bioinformatics methods, avoiding the *de novo* co-assembly process inherent to HRSM workflows. Here, we adopted an approach where we used real SM datasets from four distinct environments and sequencing depths to get insights on 1) how much data is actually needed to obtain reliable microbial ecology results and 2) the consequences on computational resources required to analyze ever expanding SM datasets - a topic often overlooked and poorly considered during planning of a shotgun metagenomic project. We performed this study using a state of the art high-resolution shotgun metagenomic methodology, which compared to other methods such as reads-based and sample-centric assembly methods (*i.e.* performing a separate de novo assembly of each sequence library) is arguably the bioinformatic method that allows the generation of the most comprehensive and meaningful end results from raw SM datasets, including the generation of beta-alpha-diversity metrics, full length genes and MAGs [2]. In that regard, the human gut microbiome dataset selected for this study (PRJNA588513) was of particular interest, as it is one of the first massive publicly available SM dataset (12,389 Gb of raw sequence data) enabled by the Illumina NovaSeq platform generated as part of a single finite project and offers a glimpse of what kind of sequencing output could be routinely achieved in terms of sequencing output in the near future for metagenomic projects. At first glance, obtaining lots of sequencing data for a SM project might seem like a good thing, but since a sequencing dataset of that magnitude cannot be readily used in its entirety in a HRSM workflow (i.e. storage of raw and intermediate files; RAM requirements for co-assembly) on the vast majority of HPC systems, how to extract the most of it has yet to be explored. In order to shed some lights on these questions, we favored an experimental design where the complete dataset was randomly subsampled at various loads to mimic various sequencing depths.

From our results with the human gut dataset, it is clear that the 0.1M clusters dataset (or 0.03 Gbases / sample; total of 24.43 Gbases) was not enough to achieve a good correlation with the results of other subsampled analyses. However, because this dataset had 912 samples, sub-sampling slightly higher, at levels as low as 0.5M clusters / sample (0.14 Gbases / sample; total of 130.25 Gbases) resulted in acceptable correlations with larger subsampled datasets. Even though the number of bases at 0.5M clusters sampling is objectively low on a per-sample basis, the corresponding total amount of bases can be considered adequate for capturing accurate population structure metrics. Moreover, even if the variability of the microbial diversity in each sample is highly dispersed (*i.e.* alpha diversity results in Fig. 3A), the fact that pooling 912 samples of 0.14 Gb each is enough to obtain a sound co-assembly and corresponding downstream results suggest that there is a redundant core of microbes common and abundant enough to many samples so that this allows for the pooling of only a tiny fraction of each library to obtain a decent quality co-assembly and consequently accurate downstream results. Pooling multiple libraries also has the advantage of putting in evidence rare reads of rare genomes that, when pooled together, reach a critical coverage level enabling their integration into the *de novo* co-assembly. The estimations of required reads for SM projects are usually considered on a per-sample basis, but this information should ideally be combined with sample evenness and the total number of samples that are part of the project. Accordingly, the number of required bases per sample should probably not be calculated on a per-sample basis, but on the total amount of bases required to obtain a reasonable co-assembly for the investigated biological system. In the case of the human gut dataset investigated here, it could be argued that 130 (0.5M clusters / sample) or 244 Gb (1M clusters / sample) of input data used for the co-assembly gave end results that were overall very similar to the results of the largest subsampled dataset of 2,927 Gb (12M clusters). In more practical terms, this suggests that this set of 912 samples could have been sequenced on one or two lanes of NovaSeq6000 S4 and give very similar results for a fraction of the cost of the original study. Similar trends were observed with the Antarctic SM dataset. In this case however, since the number of samples was much lower (*i.e.* 18) and highly variable from one another, the lowest subsampling level that gave the results mostly similar to the total dataset was approximately 20 Gbases (4M clusters / sample or 1.12 Gbases / sample). For the agricultural dataset, 0.1M clusters per sample (or 0.78 Gb) was evidently too low to obtain interpretable results. However, increasing the input to as low as 0.5M clusters / sample (3.88 Gb) enabled the generation of results highly similar to the ones obtained with the complete dataset. Based on the species discovery curve generated for our agricultural soil example dataset, (Fig. 7A), it is unlikely that sequencing saturation of the agricultural soil environment could be even remotely reached using current sequencing systems. Even if that would be possible, the amount of sequences needed would be problematic for analyses.

### The effect of community structure and richness on the required sequencing depth

How well a community is represented by a given sampling effort can be expressed in terms of the coverage of the community [15] or in terms of the coverage of the species present in the sample, *i.e.* the proportion of the overall sample richness that was detected. Besides the sampling effort, these coverage metrics are a function of the evenness and the species richness of the sample of interest. In general, the majority of a community (see 95% example in Fig. S7) can be covered by a significantly lower sampling effort than what would be necessary to cover the majority of the species richness of a sample (Fig. S8). This conclusion is very well demonstrated by our inability to exhaustively detect all species of communities of even very low species richness, like the staggered mock community used in this study, while at the same time already covering the majority of the community with very shallow subsets. As high species coverage appears to be a difficult goal to achieve, it could be a more practical alternative to calculate target sequencing depths based on a desired community coverage. This metric (community coverage) could also be used to define meaningful subsampling thresholds for already sequenced datasets.

### Compute resources

In the field of SM, the question of how many reads should be generated per sample has always been critical and the subject of continuing discussions. Currently, the most common accepted answer is probably along the lines of: the more the better - depending on the available funds of course. However, with the significant increase in sequencing throughput seen recently with the Illumina NovaSeq platform, we have reached a tipping point where computational capabilities required to perform a sound HRSM workflow are not sufficient to integrate all of the generated data anymore. In the current study it was simply not possible to perform a co-assembly of 12 Tb worth of SM data. By extrapolating from the relation between RAM as a function of input data (Table I; Fig. S1), co-assembling the human gut microbiome dataset would require approximately 10 TB of RAM and 900 hours (37.5 days) of continuous compute real time, which would translate into 57,600 core·hour (or 6.57 core·year). On most governmental and academic HPC systems, resources are usually allocated on a yearly basis and have to be carefully managed. In that regard, completely processing such a large dataset would inevitably represent a major sink in consumption of allocated resources, leaving few resources for the data processing of other projects.

### MAGs

The one area where having more data unquestionably improved end results was for MAGs generation (Fig. 6). For all SM datasets investigated here, the number and quality of MAGs increased with the number of reads used for the co-assembly. Moreover, MAGs generation was the only area where using the arm (all reads mapped) workflow undoubtedly benefited end results metrics (*i.e.* yield and quality of MAGs). There are a number of software packages to generate MAGs, here we presented results generated with MetaBAT2, essentially because the other implemented method in our workflow, Maxbin2, could not complete even after more than 200 hours of runtime on a 16 cores compute node for the human gut dataset.

### Conclusion

The main driver that prompted us to perform this *in silico* study is the foreseeable inability to use all of the reads generated by the most recent sequencing platforms for a given shotgun metagenomics project in a HRSM type of workflow. We anticipate that even our largest memory compute nodes will hardly keep up with the amount of sequences to consider all of the reads in the *de novo* assembly process inherent to HRSM workflows. Disk storage also becomes a concern as the total file size of the sequencing library files and the intermediate data that has to be transiently stored on rapid disk storage is significant. For instance, for the human gut microbiome 12M clusters arm workflow, the total compressed file size including intermediate files (*i.e.* filtered fastqs, bam, co-assembly, abundance matrices) was approximately 20 TB. Therefore, the always increasing output of sequencing data is a significant concern for the stress imposed even on large high performance computing systems. As access for HPC resources becomes increasingly competitive (see for instance 2021 Resource Allocations Competition Results), fewer resources can be allocated to an ever increasing pool of research needs. This underscores the importance of refining strategies for analyzing SM data and finding alternative advantageous ways to make good use of all the reads generated by latest sequencing technologies. In this regard, a type of workflow of potential interest would be to assemble each sample individually and combine the assembled contigs to generate a single consensual assembly. This workflow would have the advantage of eliminating the limitations in memory as de novo assembling a single library at a time should not require sizable amounts of RAM. The execution of such a workflow would however require an accepted computational method to merge or combine multiple assemblies together, which to the best of our knowledge does not exist. Software reported to perform this type of merging task are usually not maintained anymore and not suitable for modern large metagenomic assemblies [19–21] and target single genome assemblies [22–24]. At first glance, long reads (PacBio, Oxford Nanopores) can also seem attractive to replace short-reads in the objective of obtaining more contiguous co-assemblies, but performing a multi-library co-assembly of long reads data type also requires enormous amounts of RAM and compute time (personal observations), especially if reads need to be corrected prior to be assembled, as it is often the case with error-prone long reads data types.

Overall the findings reported here suggest that shallow sequencing, up to a certain level, allows reaching similar conclusions that could be reached with deep sequencing. The only exception to this is if maximizing the yield of high quality MAGs is a primary outcome or if there is a particular interest in rare functions or taxa. These conclusions are prone to have significant impacts on the planning of shotgun metagenomic projects as - in light of the results presented here - the number of sequences per sample does not need to be overly abundant to reach sound conclusions of a biological system.

## Methods

### Bioinformatics

Sequencing libraries were processed in ShotgunMG, our metagenomics bioinformatics pipeline [25–27] that was developed over the GenPipes workflow management system [28]. Sequencing adapters were first removed from each read and bases at the end of reads having a quality score <30 were cut off (Trimmomatic v0.39;[29]) and scanned for sequencing adapters contaminants reads using BBDUK (BBTools v38.1) [30] to generate quality controlled (QC) reads. The QC-passed reads from each sample were co-assembled using MEGAHIT v1.2.9 [6] on a 3 terabytes of RAM compute node with iterative kmer sizes of 31, 41, 51, 61, 71, 81, 91, 101, 111, 121 and 131 bases. MetaSPAdes [7] was also considered for co-assembly, but could not complete even after several days of computing for the lowest input human gut dataset. *Ab initio* gene prediction was performed by calling genes on each assembled contig using Prodigal v2.6.3 [31]. Assignment of KEGG orthologs (KO) was done by using DIAMOND Blastp v2.0.8 [32] to compare each predicted gene amino acids sequence of the co-assembly against the KEGG GENES database [33,34] (downloaded on 2020-03-23). COG orthologs were assigned using RPSBLAST (v2.10.1+) [35] with the CDD training sets (ftp.ncbi.nlm.nih.gov/pub/mmdb/cdd/little_endian/). The QC-passed reads were mapped (BWA mem v0.7.17; [36]) against contigs to assess quality of metagenome assembly and to obtain contig abundance profiles. Alignment files in bam format were sorted by read coordinates using samtools v1.9 [37], and only properly aligned read pairs were kept for downstream steps. Each bam file (containing properly aligned paired-reads only) was analyzed for coverage of contigs and predicted genes using bedtools (v2.23.0; [38]) using a custom bed file representing gene coordinates on each contig. Only paired reads both overlapping their contig or gene were considered for gene counts. Coverage profiles of each sample were merged to generate an abundance matrix (rows = contig, columns = samples). Taxonomic lineage assignment to each contig was performed using CAT v5.2.3 [39] with the following key parameters (*f*=0.5; *r*=1). Taxonomic summaries and beta diversity metrics were computed with microbiomeutils v0.9.4 [40]. Alpha diversity metrics were obtained with RTK v0.93.2 [41]. MAGs were generated using MetaBAT (v2.12.1) [42] using an abundance matrix generated with the jgi_summarize_bam_contig_depths software [42] with the — minContigLength 1000 —minContigDepth 2 and —minContigIdentity 97 parameters. The quality of obtained MAGs was assessed with CheckM v1.1.3 [43]. MaxBin2 [44] was also considered, but could not be completed even after several days of computing the lowest input human gut dataset.

### rpoB-based analyses and simulations

Genes encoding the β subunit of the RNA polymerase (*rpoB*), a single-copy gene universally present in microorganisms, were identified based on genes assigned to the COG [45] definition “COG0085”. Counts of reads mapped to these genes were treated as gene abundances and were summarized as sample-specific *rpoB* gene count tables. The true sequencing depth of samples was calculated based on the sequencing depth of the subset corrected by sample purity. Sample purity was calculated as the ratio of reads mapped to contigs classified as being of archaeal or bacterial origin over the sum of all mapped reads. Pileou’s evenness index was calculated based on the *rpoB* gene count tables and from the results of the specnumber and diversity functions of the R package “vegan” [46]. The species coverage of data subsets was calculated on a per-sample basis, using the richness of the complete dataset (agricultural soil, Antarctic) or the 12M subset (human gut) as a reference. As genes were not linked between subsets, community coverage [15] was estimated by interpolation using the Coverage function of the R package “entropart” [47]. In detail, *rpoB* count tables of the complete (agricultural soil, Antarctic) or the 12M subset (human gut) datasets were passed to the function and the community coverage was interpolated for the *rpoB* reads count levels that were obtained for the individual subsets. Consistent with the species coverage metric, community coverage was estimated on a per-sample basis.

The effects of adding additional samples while maintaining a constant sequencing depth were investigated based on the obtained *rpoB* count tables and using a custom R script. In detail, a reference meta-community for each dataset was calculated by adding up all *rpoB* count tables (rarefied to a common depth; rrarefy function; [46] of the samples of this dataset. The overall species richness and the mean community structure of this meta-community were then defined as the reference for each dataset. The rarefied *rpoB* count tables were subsequently combined in combinations of 2 to n_max_ samples, where n_max_ represented the total number of samples in the dataset; i.e. the script iterated through C(n_max_, 2) to C(n_max_,n_max_). Due to the immense computational burden of exhaustively simulating all possible combinations, only n_max_ random combinations for each combination level were calculated. All count tables were rarefied to a constant depth after combination. The corresponding species coverage was calculated as the proportion of the detected species of a sample combination over the overall species richness of the meta-community. The corresponding community coverage of the combined *rpoB* count tables was calculated as the sum of the relative abundances of the species’ of the meta-community that were detected in the community of the combination. The R script to perform this simulation is provided as part of this study.

### Functional analyses

For each predicted gene, the best hit having at least an e-value ≤ 1e-10 against the KEGG genes database was kept as the KEGG representative of that gene. Similarly, COG representative of each gene corresponded to the best hit COG hit having at least an e-value ≤ 1e-10. Each assigned KEGG gene may be associated with a KEGG ortholog (KO). For the analyses of figures 4 and S4, read counts all the genes from the human gut or antarctic gene abundance matrix that were assigned with the same KO were summed to generate a KO abundance matrix. For each library, each KO aggregated value was divided by the total read abundance of the *recA* gene (COG0468) to obtain a normalized value for each KO.

Statistics and figures were generated using R v4.1.2 for which the code is available on github (https://github.com/jtremblay/shotgunmg_paper).

### Availability of source code and requirements

ShotgunMG - http://jtremblay.github.io/shotgunmg.html

<u>The ShotgunMG pipeline wrapper code and Python, Perl and R scripts that are being called by ShotgunMG are available here</u>: https://bitbucket.org/jtremblay514/nrc_pipeline_public/src/1.3.0/

https://bitbucket.org/jtremblay514/nrc_tools_public/src/1.3.0/

<u>External software packages module install scripts are available here:</u> https://bitbucket.org/jtremblay514/nrc_resources_public/src/1.3.0/

A Docker image built on the CentOS 7 operational system which contains all necessary modules for full pipeline functionality is available for testing/evaluation purposes and running small datasets (https://cloud.docker.com/u/julio514/repository/docker/julio514/centos).

### Availability of supporting data and materials

Human gut microbiome dataset: PRJNA588513

Antarctic soil dataset: PRJNA513362

Agricultural soil dataset: PRJNA513362

Mock communities: PRJNA873699

An example of the complete list of commands with all parameters for each package used in our analysis pipeline for the human gut, Antarctic, agricultural soil and mock communities datasets and all the key results generated during this study, including de novo assemblies, genes and contigs abundance matrices, functional and taxonomic annotations and MAGs are available on zenodo.org: https://doi.org/10.5281/zenodo.6349279.

## Declarations

### List of abbreviations

Tb: Tera-base
TB: Terabyte
Gb: Giga-base
GB: Gigabyte
HPC: High Performance Computing
bp: base-pairs
MB: Megabyte
SM: Shotgun metagenomics
SSM: Shallow Shotgun Metagenomics
HRSMG: High resolution shotgun metagenomics
KO: KEGG ortholog
arm: all reads mapped.

## Author contributions

JT planned the experimental design, wrote the software, analyzed the data and wrote the manuscript. LS contributed the *rpoB*-based analyses and wrote the corresponding part of the manuscript. CWG analyzed results and edited the manuscript.

Julien Tremblay is a Senior Research Officer at the National Research Council of Canada. His research focuses on end-to-end bioinformatics pipelines architecture and development geared for metagenomics data and using sequence-based approaches to understand microbial communities.

Lars Schreiber is an Associate Research Officer at the National Research Council of Canada. His research focuses on understanding how microbial communities interact with their environment and how this interaction is affected by stressors.

Charles W Greer is a Principal Research Officer and the Group Leader of the Genomics and Microbiomes Group at the National Research Council of Canada. His research interest is to monitor microbial community structural and functional diversity related to ecosystem processes using metagenomics and metatranscriptomics.

## Acknowledgments

We wish to acknowledge Compute Canada for access to both the Waterloo University (Graham system) infrastructure. We acknowledge Shared Services Canada for access to the General Purpose Scientific Cluster (GPSC).

---

**Figure S1.** Statistics on the various co-assemblies generated.

**Figure S2.** Statistics of aligned reads on co-assembly references for the A) human gut, B) Antarctic, C) agricultural soil and D) mock communities datasets. arm = all reads mapped workflow.

**Figure S3.** PCoA plots of beta-diversity of each dataset at each sequencing depth. For the A) human gut dataset, samples were found to cluster according to their diversity level (i.e. diversity quantiles) with high diversity individuals clustering together and low diversity individuals clustering together as well. Samples were colored by sampling location for the B) Antarctic dataset and followed a similar structure throughout the various sequencing depths. C) Agricultural soil samples clustered according to their treatment in nitrogen and D) mock communities according to their spiked-in configuration.

**Figure S4.** Taxonomic coverage analyses for the A) human gut, B) Antarctic, C) agricultural soil and D) mock communities datasets. Top left panel: Spearman correlation coefficients of Bray-Curtis dissimilarity matrices computed from relative abundance taxonomic summary tables of each workflow. The Higher coefficient means higher correlation between taxonomic summaries. Top right panel: Number of unique taxonomic lineages observed as a function of each workflow’s input data load. Lower left panel: Venn diagram of each unique taxonomic lineage as a function of data input load workflows. Lower right panel: Number of reads associated with each category listed in the Venn diagram.

**Figure S5.** KEGG orthologs (KO) coverage analyses for the A) human gut, B) Antarctic, C) agricultural soil and D) mock communities datasets. Top left panel: Spearman correlation coefficients of Bray-Curtis dissimilarity matrices computed from normalized KO abundance tables of each workflow. The Higher coefficient means higher correlation between KO abundance profiles. Top right panel: Number of unique KO observed as a function of each workflow’s input data load. Lower left panel: Venn diagram of each unique KO as a function of data input load workflows. Lower right panel: Number of reads associated with each category listed in the Venn diagram.

**Figure S6.** Independent in silico repetition of the standard workflow for the human gut dataset for 0.1M, 0.5M and 1M sequencing clusters per sample.

**Figure S7.** The effects of sampling effort and community richness and evenness on the achieved community coverage. The shown surface represents the required sampling effort to achieve 95% community coverage for communities of varying species richness and evenness J. Sampling effort is additionally also indicated by surface colour. The data for this figure is based on randomly drawing members of simulated communities with varying evenness and richness.

**Figure S8.** The effects of sampling effort and community richness and evenness on the achieved species coverage. The shown surface represents the required sampling effort to achieve 95% species (richness) coverage for communities of varying species richness and evenness J. Sampling effort is additionally also indicated by surface colour. The data for this figure is based on randomly drawing members of simulated communities with varying evenness and richness.

## References

1. Breitwieser FP, Lu J, Salzberg SL. A review of methods and databases for metagenomic classification and assembly. Brief. Bioinform. 2019; 20:1125–1136

2. Chao Yang, Debajyoti Chowdhury, Zhenmiao Zhang, William K. Cheung, Aiping Lu, Zhaoxiang Bian, Lu Zhang. A review of computational tools for generating metagenome-assembled genomes from metagenomic sequencing data. Comput. Struct. Biotechnol. J. 2021; 19:6301–6314

3. Georganas E, Egan R, Hofmeyr S, et al. Extreme Scale De Novo Metagenome Assembly. SC18: International Conference for High Performance Computing, Networking, Storage and Analysis 2018;

4. Boisvert S, Raymond F, Godzaridis E, et al. Ray Meta: scalable de novo metagenome assembly and profiling. Genome Biol. 2012; 13:R122

5. Sczyrba A, Hofmann P, Belmann P, et al. Critical Assessment of Metagenome Interpretation-a benchmark of metagenomics software. Nat. Methods 2017; 14:1063–1071

6. Li D, Liu C-M, Luo R, et al. MEGAHIT: an ultra-fast single-node solution for large and complex metagenomics assembly via succinct de Bruijn graph. Bioinformatics 2015; 31:1674–1676

7. Nurk S, Meleshko D, Korobeynikov A, et al. metaSPAdes: a new versatile metagenomic assembler. Genome Res. 2017; 27:824–834

8. Meyer F, Fritz A, Deng Z-L, et al. Critical Assessment of Metagenome Interpretation - the second round of challenges. bioRxiv 2021;

9. Coleine C, Albanese D, Onofri S, et al. Metagenomes in the Borderline Ecosystems of the Antarctic Cryptoendolithic Communities. Microbiol Resour Announc 2020; 9:

10. Sun Y, Zuo T, Cheung CP, et al. Population-Level Configurations of Gut Mycobiome Across 6 Ethnicities in Urban and Rural China. Gastroenterology 2021; 160:272–286.e11

11. Li Y, Tremblay J, Bainard LD, et al. Long-term effects of nitrogen and phosphorus fertilization on soil microbial community structure and function under continuous wheat production. Environ. Microbiol. 2020; 22:1066–1088

12. Tremblay J, Greer CW. Shotgun metagenomic sequencing dataset of a synthetic mock community containing 20 genomes spiked-in at even and staggered concentrations. Submitted

13. Case RJ, Boucher Y, Dahllöf I, et al. Use of 16S rRNA and rpoB genes as molecular markers for microbial ecology studies. Appl. Environ. Microbiol. 2007; 73:278–288

14. Pielou EC. The measurement of diversity in different types of biological collections. J. Theor. Biol. 1966; 13:131–144

15. Chao A, Jost L. Coverage-based rarefaction and extrapolation: standardizing samples by completeness rather than size. Ecology 2012; 93:2533–2547

16. Hillmann B, Al-Ghalith GA, Shields-Cutler RR, et al. Evaluating the Information Content of Shallow Shotgun Metagenomics. mSystems 2018;

17. Xu W, Chen T, Pei Y, et al. Characterization of Shallow Whole-Metagenome Shotgun Sequencing as a High-Accuracy and Low-Cost Method by Complicated Mock Microbiomes. Front. Microbiol. 2021; 0:

18. Snipen L, Angell I-L, Rognes T, et al. Reduced metagenome sequencing for strain-resolution taxonomic profiles. Microbiome 2021; 9:1–19

19. Scholz M, Lo C-C, Chain PSG. Improved assemblies using a source-agnostic pipeline for MetaGenomic Assembly by Merging (MeGAMerge) of contigs. Sci. Rep. 2014; 4:6480

20. Vicedomini R, Vezzi F, Scalabrin S, et al. GAM-NGS: genomic assemblies merger for next generation sequencing. BMC Bioinformatics 2013; 14 Suppl 7:S6

21. Soto-Jimenez L, Estrada K, Sanchez-Flores A. GARM: Genome Assembly, Reconciliation and Merging Pipeline. Current Topics in Medicinal Chemistry 2014; 14:418–424

22. Tang L, Li M, Wu F-X, et al. MAC: Merging Assemblies by Using Adjacency Algebraic Model and Classification. Front. Genet. 2020; 0:

23. Wences AH, Schatz MC. Metassembler: merging and optimizing de novo genome assemblies. Genome Biol. 2015; 16:207

24. Lin S-H, Liao Y-C. CISA: contig integrator for sequence assembly of bacterial genomes. PLoS One 2013; 8:e60843

25. Liu J, Cade-Menun BJ, Yang J, et al. Long-Term Land Use Affects Phosphorus Speciation and the Composition of Phosphorus Cycling Genes in Agricultural Soils. Front. Microbiol. 2018; 0:

26. Tremblay J, Yergeau E, Fortin N, et al. Chemical dispersants enhance the activity of oil- and gas condensate-degrading marine bacteria. ISME J. 2017; 11:2793–2808

27. . Bitbucket.

28. Bourgey M, Dali R, Eveleigh R, et al. GenPipes: an open-source framework for distributed and scalable genomic analyses. Gigascience 2019; 8:

29. Bolger AM, Lohse M, Usadel B. Trimmomatic: a flexible trimmer for Illumina sequence data. Bioinformatics 2014; 30:2114

30. Bushnell B. BBMap. SourceForge

31. Hyatt D, Chen G-L, LoCascio PF, et al. Prodigal: prokaryotic gene recognition and translation initiation site identification. BMC Bioinformatics 2010; 11:119

32. Buchfink B, Reuter K, Drost H-G. Sensitive protein alignments at tree-of-life scale using DIAMOND. Nat. Methods 2021; 18:366–368

33. Kanehisa M, Goto S. KEGG: Kyoto Encyclopedia of Genes and Genomes. Nucleic Acids Res. 2000; 28:27–30

34. Kanehisa M. Toward understanding the origin and evolution of cellular organisms. Protein Sci. 2019; 28:

35. Camacho C, Coulouris G, Avagyan V, et al. BLAST+: architecture and applications. BMC Bioinformatics 2009; 10:421

36. Li H. Aligning sequence reads, clone sequences and assembly contigs with BWA-MEM. 2013;

37. Li H, Handsaker B, Wysoker A, et al. The Sequence Alignment/Map format and SAMtools. Bioinformatics 2009; 25:2078–2079

38. Quinlan AR, Hall IM. BEDTools: a flexible suite of utilities for comparing genomic features. Bioinformatics 2010; 26:841–842

39. von Meijenfeldt FAB, Arkhipova K, Cambuy DD, et al. Robust taxonomic classification of uncharted microbial sequences and bins with CAT and BAT. Genome Biol. 2019; 20:1–14

40. Tremblay J. microbiomeutils: Python utility to generate distance matrices, perform PCoAs and generate taxonomic summaries using simple tab-separated feature tables.

41. Saary P, Forslund K, Bork P, et al. RTK: efficient rarefaction analysis of large datasets. Bioinformatics 2017; 33:2594–2595

42. Kang DD, Froula J, Egan R, et al. MetaBAT, an efficient tool for accurately reconstructing single genomes from complex microbial communities. PeerJ 2015; 3:e1165

43. Parks DH, Imelfort M, Skennerton CT, et al. CheckM: assessing the quality of microbial genomes recovered from isolates, single cells, and metagenomes. Genome Res. 2015; 25:1043

44. Wu Y-W, Simmons BA, Singer SW. MaxBin 2.0: an automated binning algorithm to recover genomes from multiple metagenomic datasets. Bioinformatics 2015; 32:605–607

45. Galperin MY, Makarova KS, Wolf YI, et al. Expanded microbial genome coverage and improved protein family annotation in the COG database. Nucleic Acids Res. 2015; 43:D261–9

46. Oksanen J, Blanchet FG, Kindt R, et al. vegan: community ecology package. version 1.17–2. 2010;

47. Marcon E, Hérault B. entropart: AnRPackage to Measure and Partition Diversity. J. Stat. Softw. 2015; 67:

