## Supplementary figures and images for "High resolution shotgun metagenomics: the more data, the better?"

### Figure S1

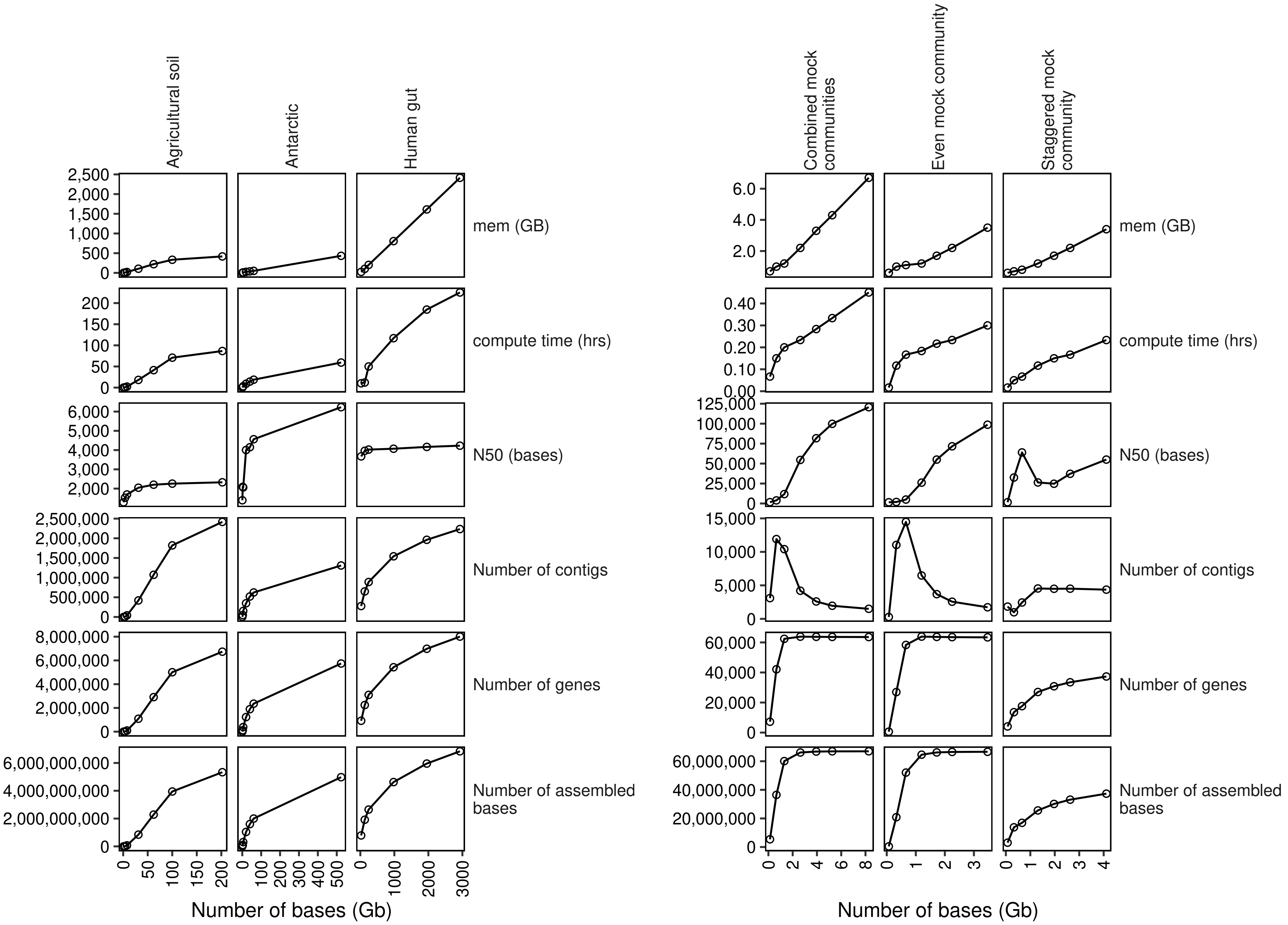

### Figure S2

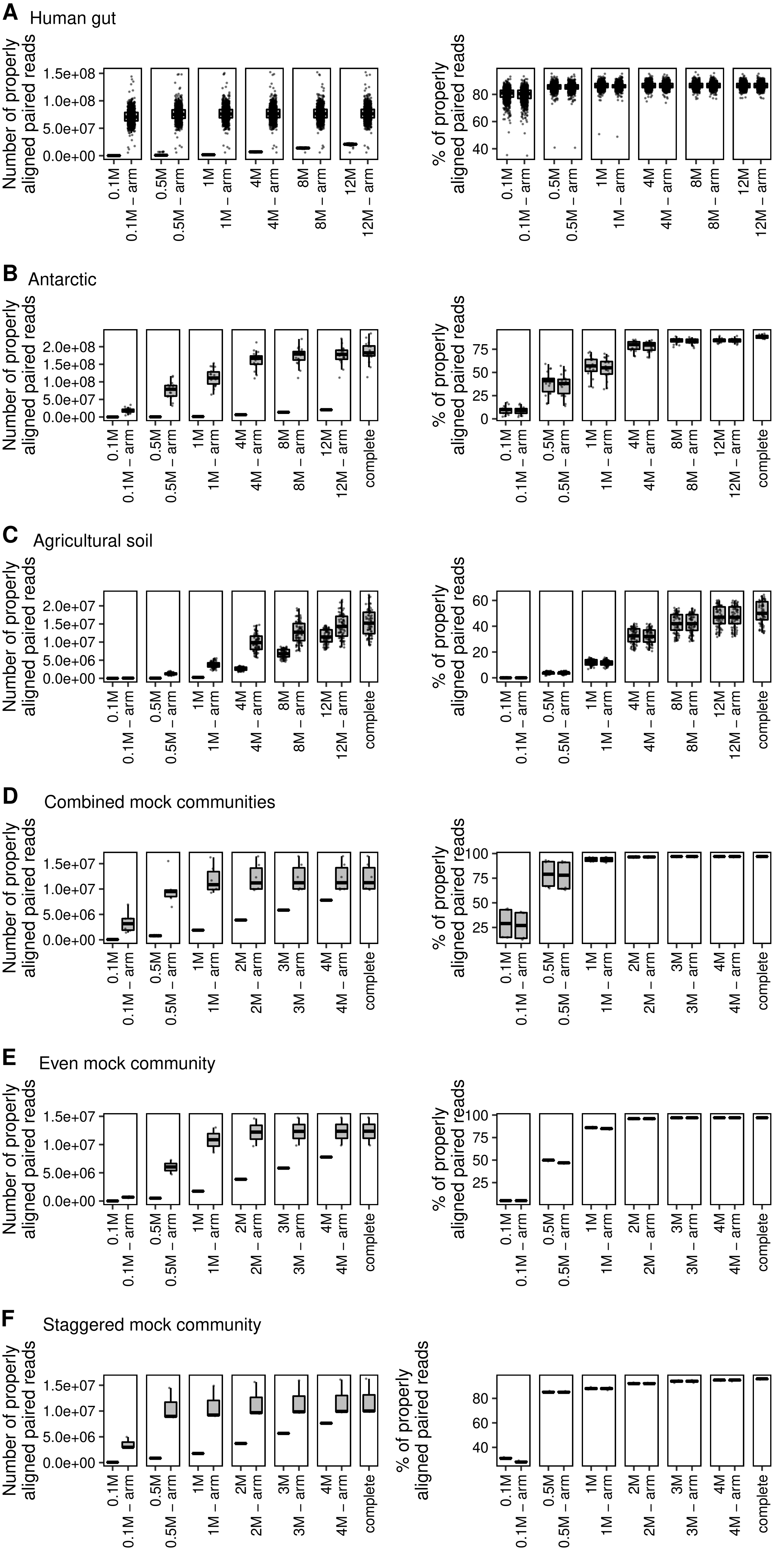

### Figure S3

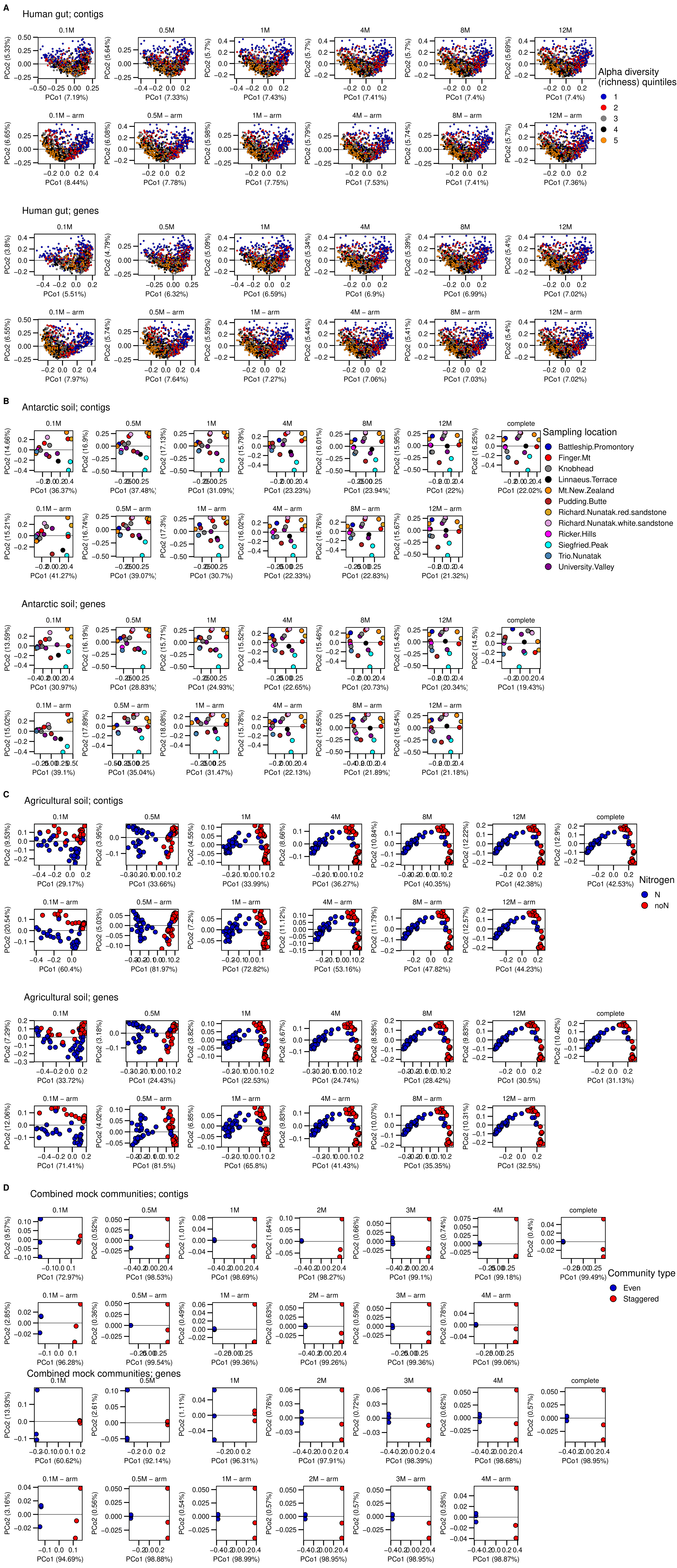

### Figure S6

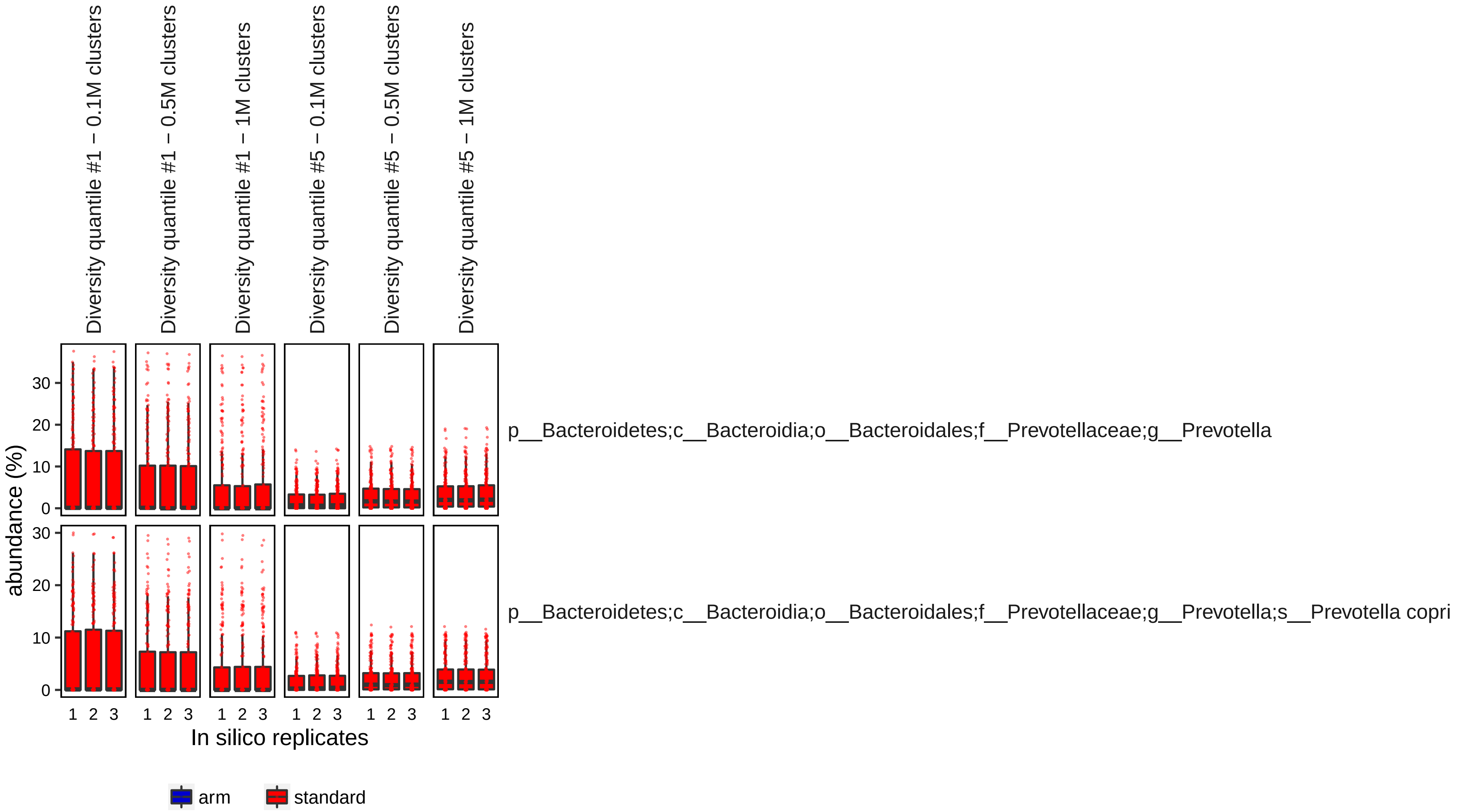

### Figure S7

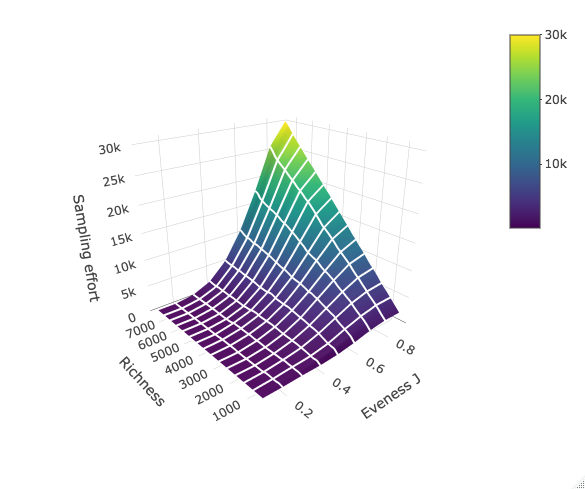

### Figure S8

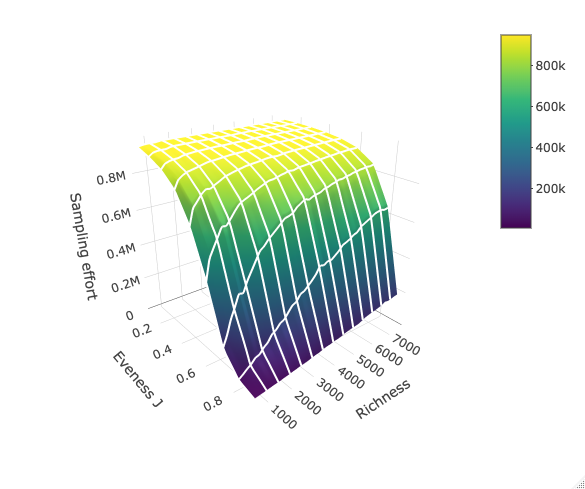
