## Supplementary material for "High resolution shotgun metagenomics: the more data, the better?": Figure S4

A

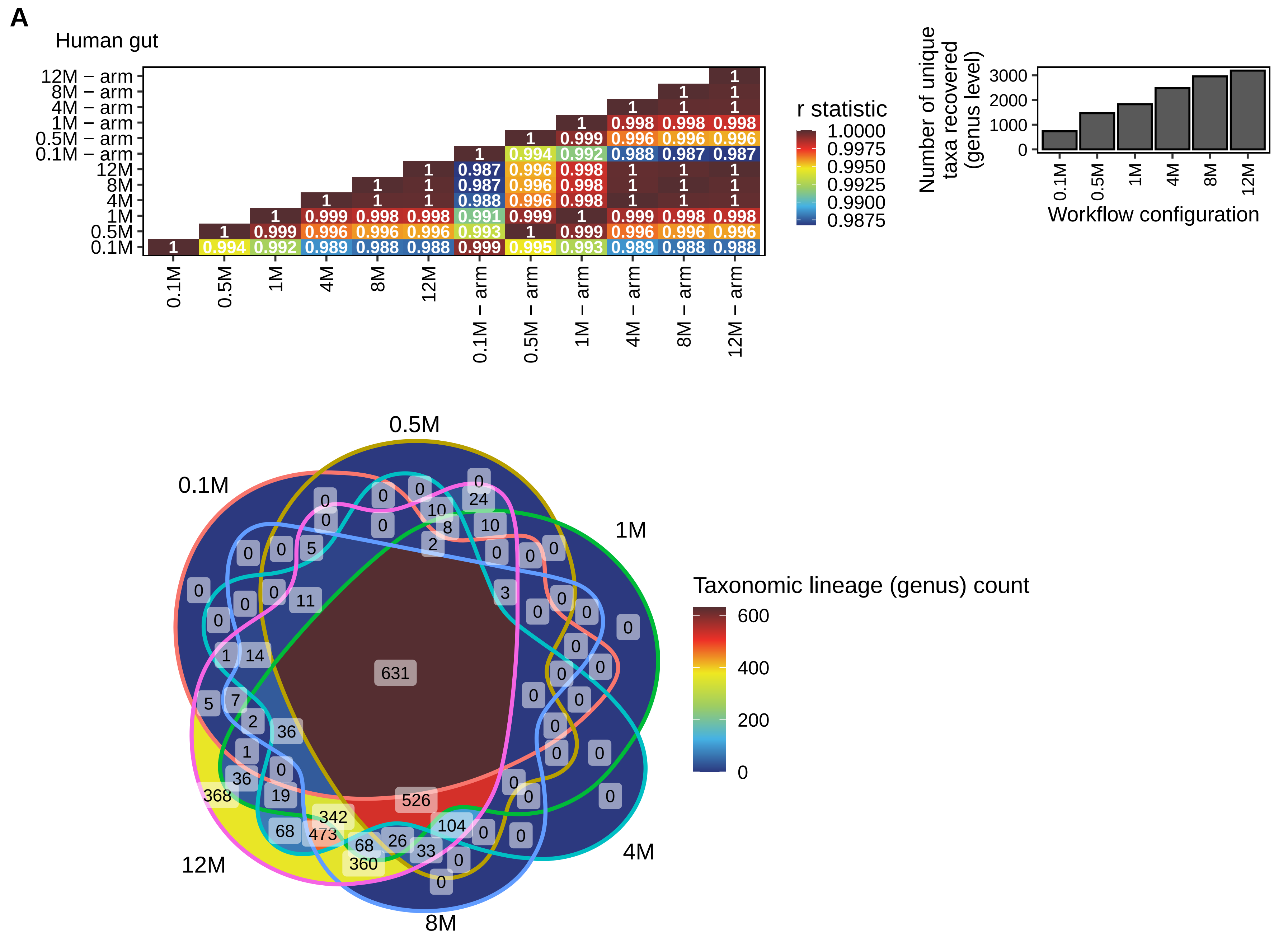


Distribution of reads for the 12M reads configuration

| | Number of reads | % |
|---|---|---|
| *A − 368* | 233,416 | 0.001 |
| *B − 473* | 913,590 | 0.005 |
| *C − 342* | 3,887,226 | 0.021 |
| *D − 360* | 185,644 | 0.001 |
| *E − 526* | 29,830,578 | 0.157 |
| *F − 631* | 18,924,535,256 | 99.815 |

A,B,C,D,E,F letters correspond to the Venn diagram zones.

B

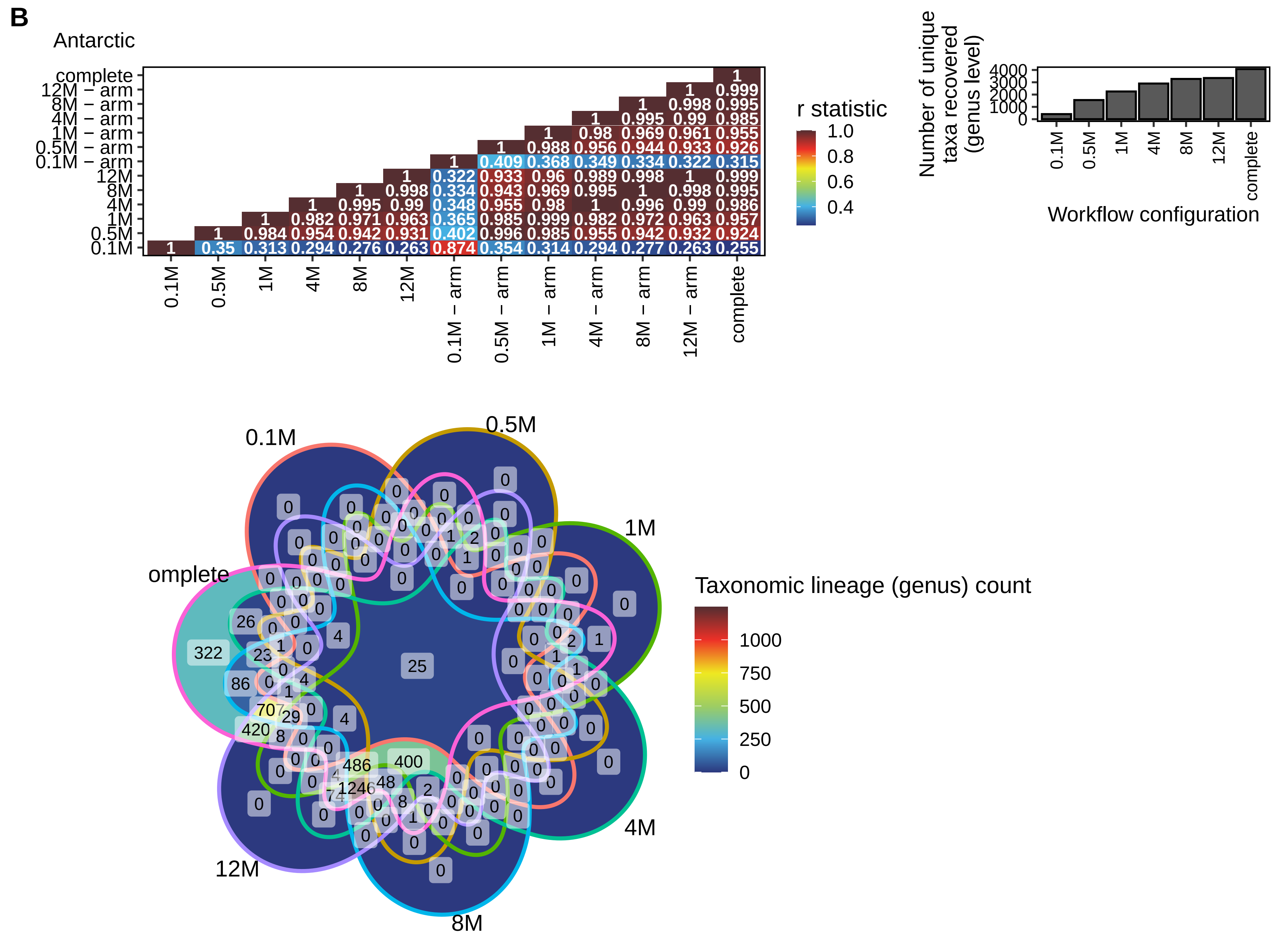


Distribution of reads for the complete dataset configuration

| | Number of reads | % |
|---|---|---|
| *A − 322* | 27,602 | 0.016 |
| *B − 86* | 13,434 | 0.008 |
| *C − 707* | 312,196 | 0.186 |
| *D − 420* | 64,320 | 0.038 |
| *E − 486* | 18,225,106 | 10.859 |
| *F − 1246* | 4,866,268 | 2.899 |
| *G − 400* | 144,330,002 | 85.993 |

A,B,C,D,E,F letters correspond to the Venn diagram zones.

C

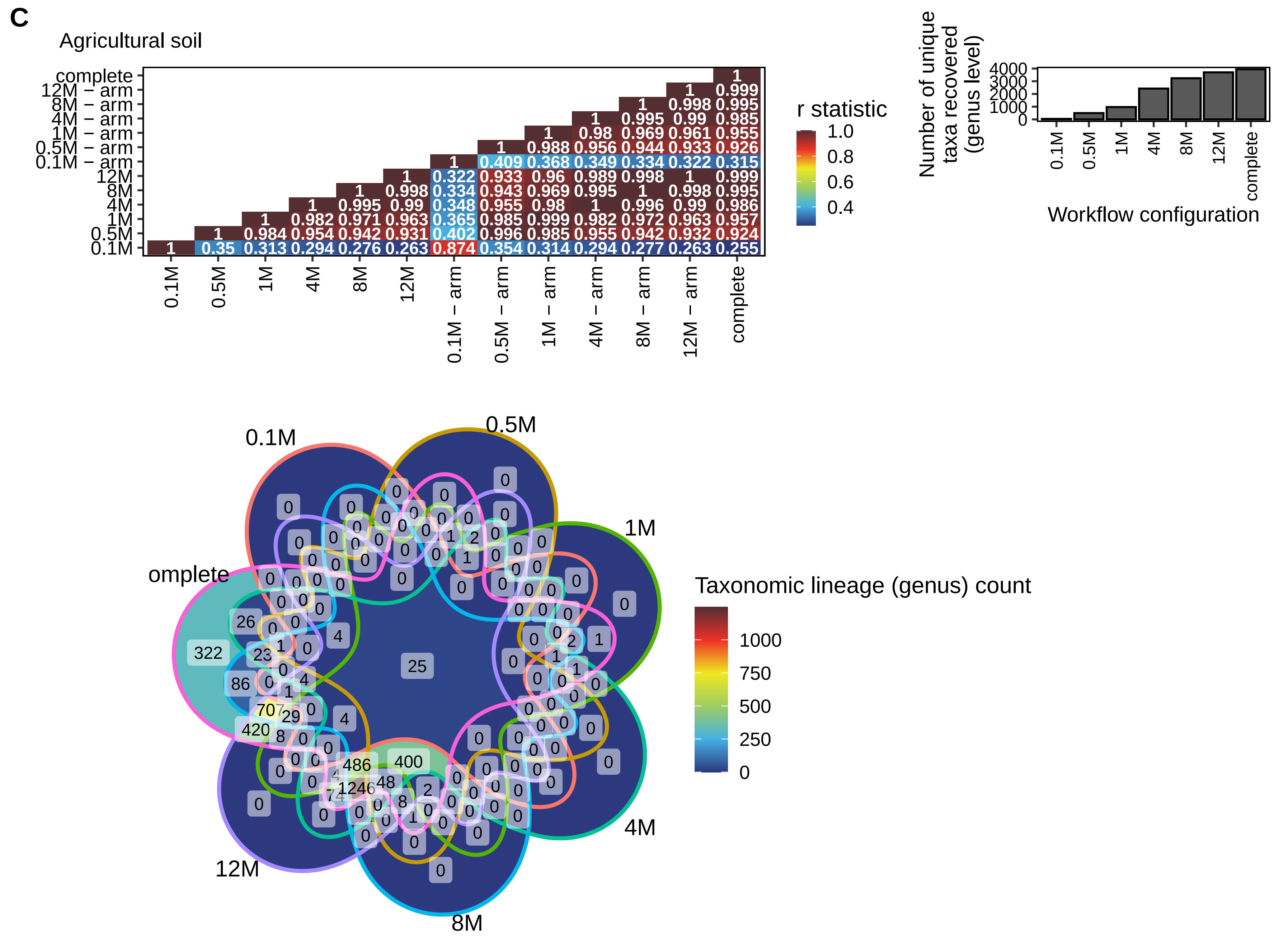


Distribution of reads for the complete dataset configuration

| | Number of reads | % |
|---|---|---|
| *A − 322* | 97,386 | 0.009 |
| *B − 86* | 52,192 | 0.005 |
| *C − 707* | 1,146,070 | 0.104 |
| *D − 420* | 237,200 | 0.021 |
| *E − 486* | 67,740,330 | 6.130 |
| *F − 1246* | 18,196,584 | 1.647 |
| *G − 400* | 540,317,808 | 48.892 |
| *H − 25* | 477,329,266 | 43.193 |

A,B,C,D,E,F,G,H letters correspond to the Venn diagram zones.

D

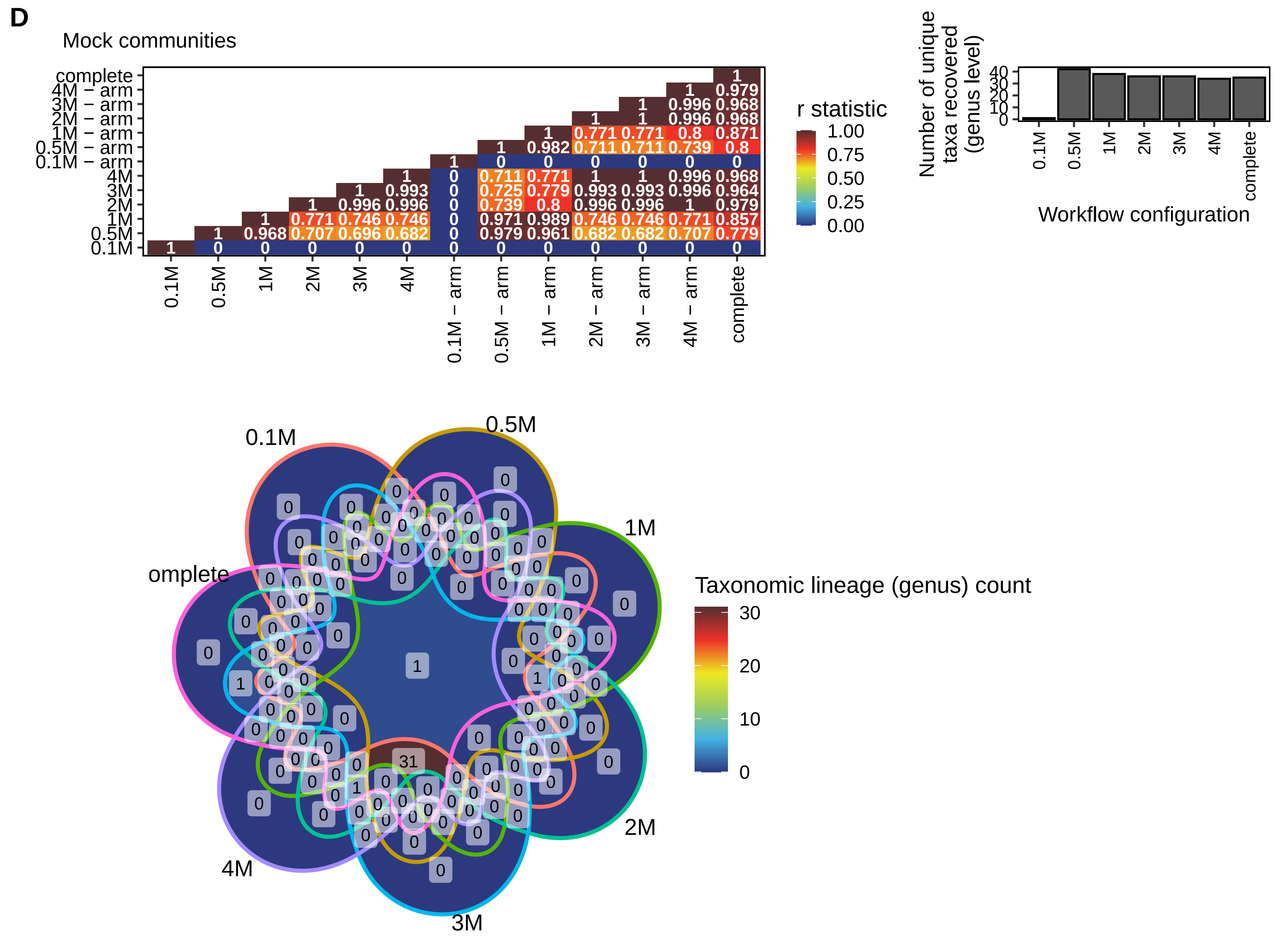


Distribution of reads for the complete dataset configurati

| | Number of reads | % |
|---|---|---|
| *A − 31* | 73,236,620 | 99.44 |
| *B − 1* | 11,344 | 0.02 |
| *C − 1* | 12,228 | 0.02 |
| *D − 1* | 389,734 | 0.53 |

A,B,C,D,E,F letters correspond to the Venn diagram zones.
