## Supplementary material for "High resolution shotgun metagenomics: the more data, the better?": Figure S5

A

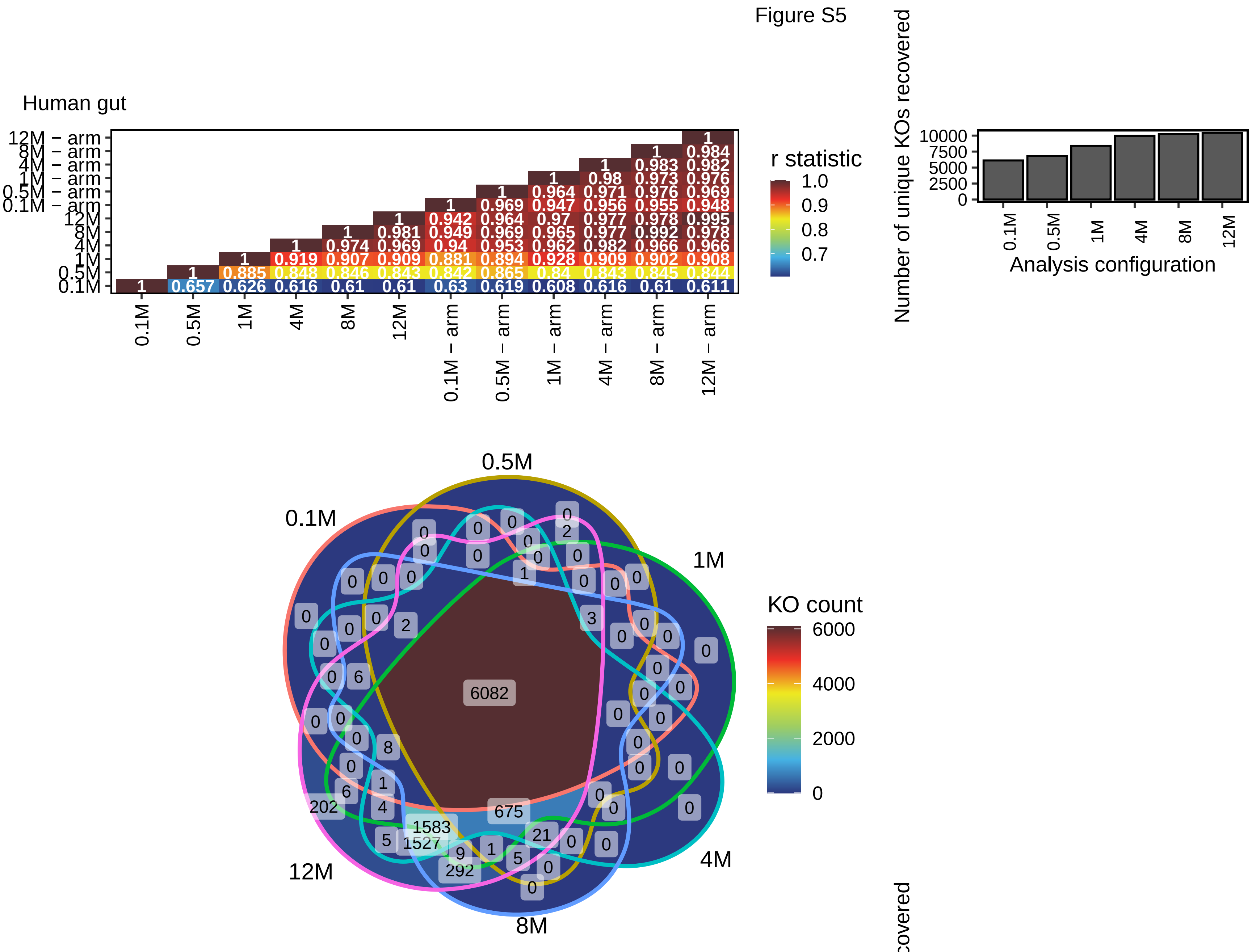


Distribution of reads for the 12M reads configuration

| | Number of reads | % |
|---|---|---|
| *A − 202* | 19,655 | 0.0002 |
| *B − 1527* | 800,906 | 0.0074 |
| *C − 1583* | 1,567,161 | 0.0145 |
| *D − 675* | 3,429,827 | 0.0317 |
| *E − 292* | 40,898 | 0.0004 |
| *F − 6082* | 10,819,128,750 | 99.9459 |

A,B,C,D,E,F letters correspond to the Venn diagram zones.

B

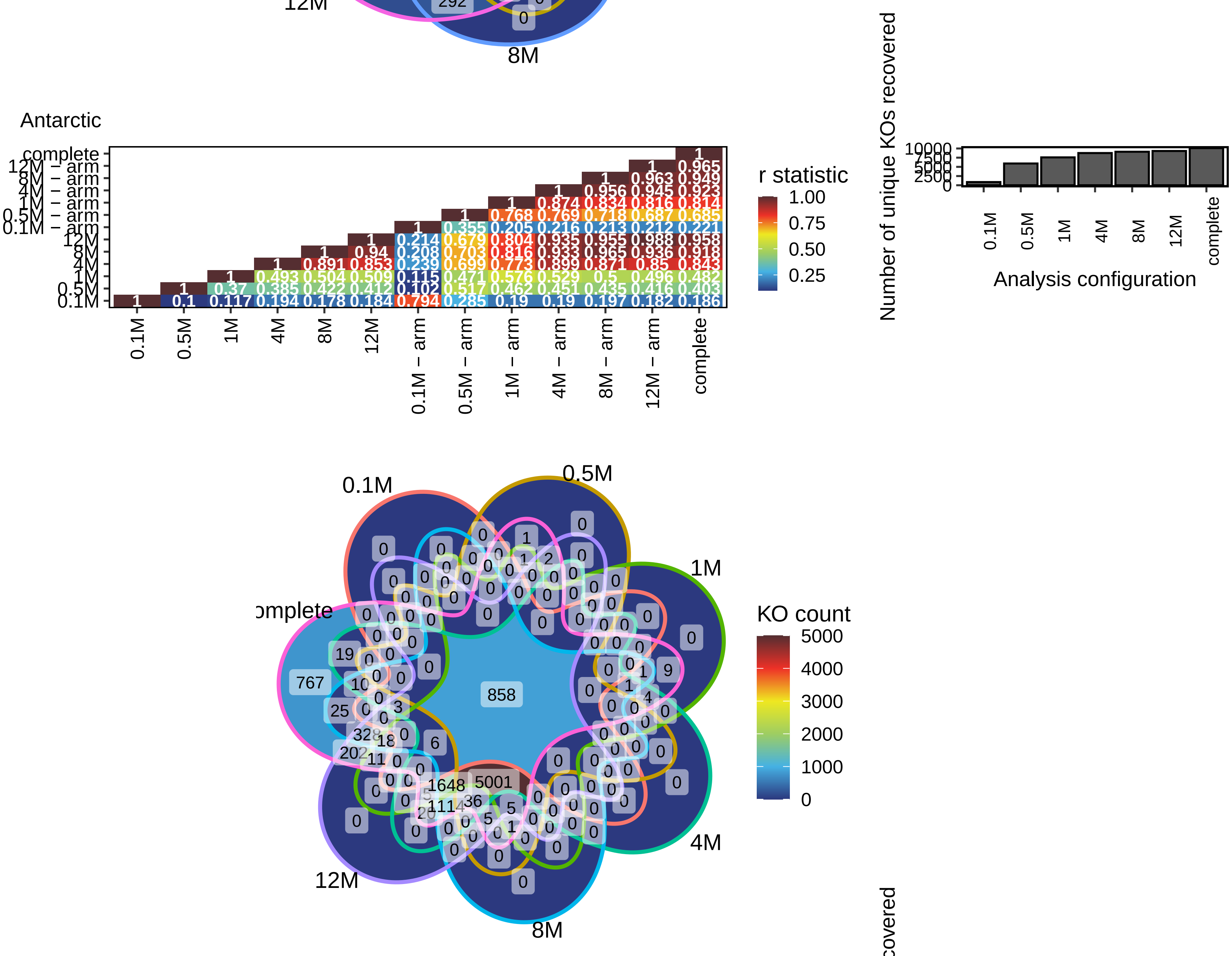


Distribution of reads for the 'complete' reads configuration

| | Number of reads | % |
|---|---|---|
| *A − 767* | 159,619 | 0.02 |
| *B − 328* | 528,266 | 0.05 |
| *C − 202* | 144,691 | 0.01 |
| *D − 1648* | 29,202,488 | 2.93 |
| *E − 1114* | 5,978,756 | 0.60 |
| *F − 5001* | 624,712,435 | 62.60 |
| *G − 858* | 337,290,304 | 33.80 |

A,B,C,D,E,F,G letters correspond to the Venn diagram zones.

C

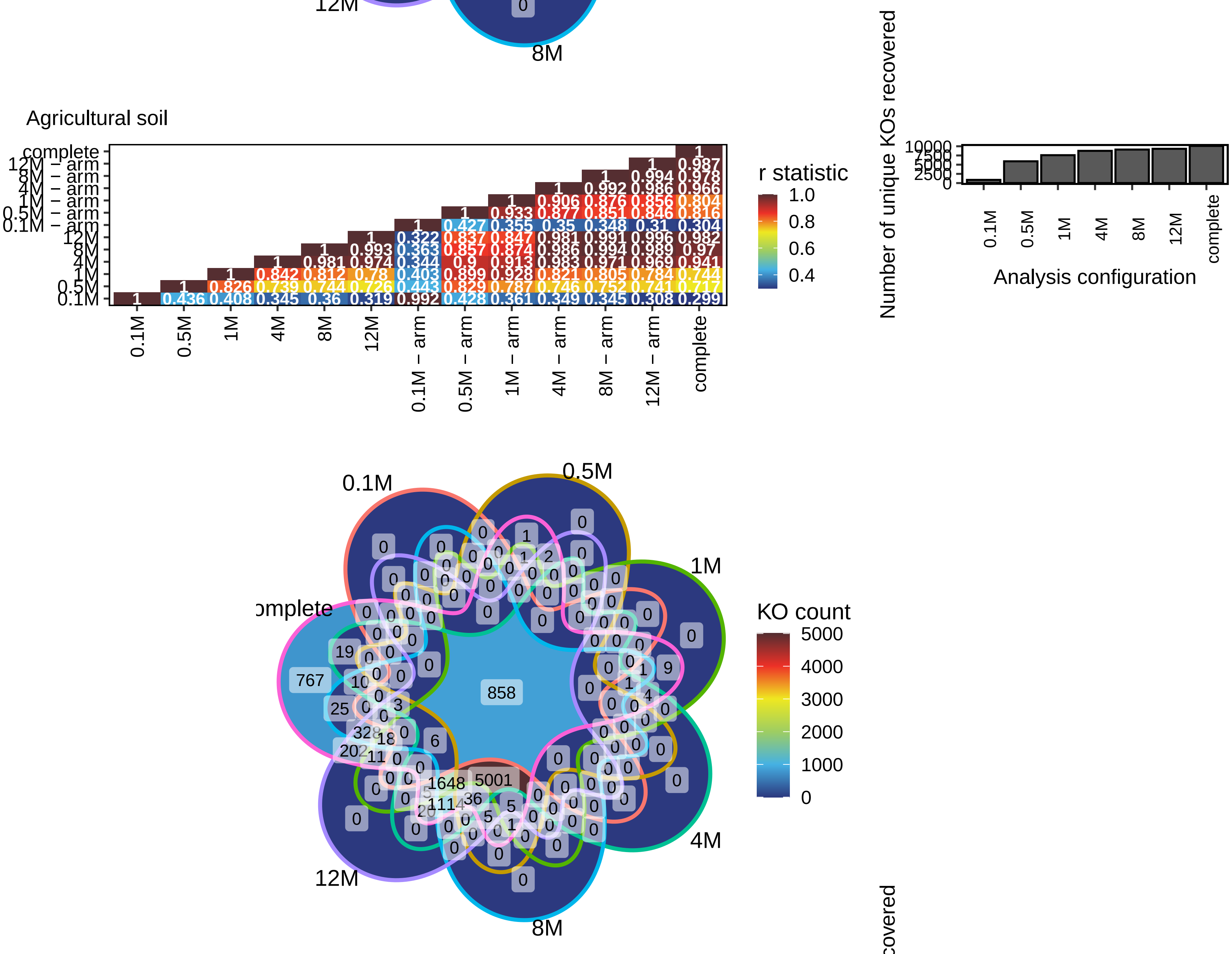


Distribution of reads for the 'complete' reads configuration

| | Number of reads | % |
|---|---|---|
| *A − 858* | 161,844,970 | 29.27 |
| *B − 328* | 1,925,564 | 0.35 |
| *C − 202* | 1,110,543 | 0.20 |
| *D − 1648* | 25,244,818 | 4.57 |
| *E − 1114* | 9,860,882 | 1.78 |
| *F − 5001* | 350,613,091 | 63.41 |
| *G − 25* | 97,281 | 0.02 |
| *H − 767* | 2,193,901 | 0.40 |

A,B,C,D,E,F,G letters correspond to the Venn diagram zones.

D

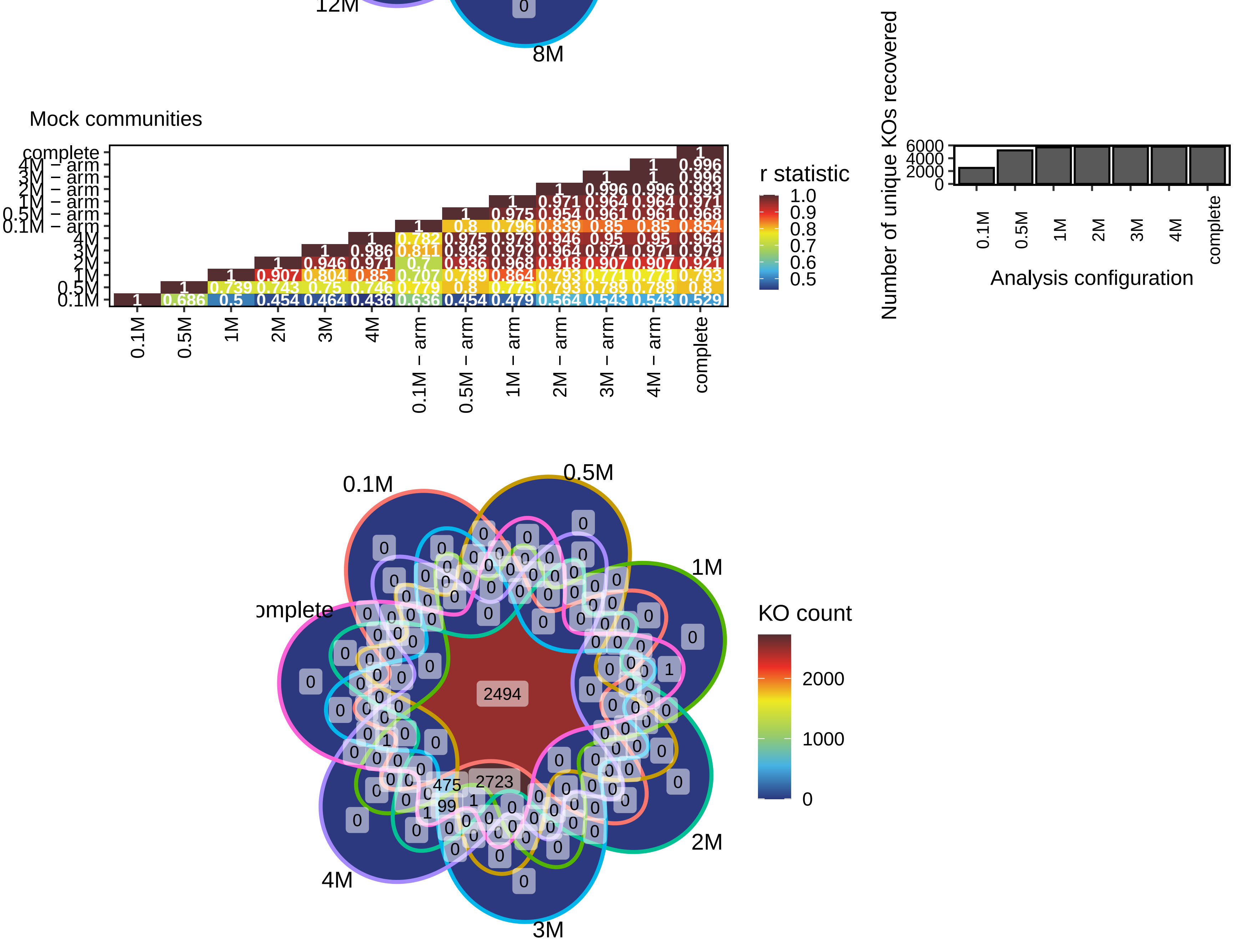


Distribution of reads for the 'complete' reads configuration

| | Number of reads | % |
|---|---|---|
| *A − 2494* | 35,944,804 | 77.8495 |
| *B − 2723* | 9,855,476 | 21.3451 |
| *C − 475* | 339,024 | 0.7343 |
| *D − 99* | 31,560 | 0.0684 |
| *E − 1* | 684 | 0.0015 |
| *F − 1* | 399 | 0.0009 |
| *G − 1* | 208 | 0.0005 |

A,B,C,D,E,F,G letters correspond to the Venn diagram zones.
